# The Heritability of Pathogen Traits - Definitions and Estimators

**DOI:** 10.1101/058503

**Authors:** Venelin Mitov, Tanja Stadler

## Abstract

Pathogen traits, such as the virulence of an infection, can vary significantly between patients. A major challenge is to measure the extent to which genetic differences between infecting strains explain the observed variation of the trait. This is quantified by the trait’s broad-sense heritability, *H*^2^. A recent discrepancy between estimates of the heritability of HIV-virulence has opened a debate on the estimators’ accuracy. Here, we show that the discrepancy originates from model limitations and important lifecycle differences between sexually reproducing organisms and transmittable pathogens. In particular, current quantitative genetics methods, such as donor-recipient regression (DR) of surveyed serodiscordant couples and the phylogenetic mixed model (PMM), are prone to underestimate *H*^2^, because they fail to model the gradual loss of phenotypic resemblance between transmission-related patients in the presence of within-host evolution. We explore two approaches correcting these errors: ANOVA on closest phylogenetic pairs (ANOVA-CPP) and the phylogenetic Ornstein-Uhlenbeck mixed model (POUMM). Empirical analyses reveal that at least 25% of the variation in HIV-virulence is explained by the virus genome both for European and African data. These results confirm the presence of significant factors for HIV virulence in the viral genotype and reject previous hypotheses of negligible viral influence. Beyond HIV, ANOVA-CPP is ideal for slowly evolving protozoa, bacteria and DNA-viruses, while POUMM suits rapidly mutating RNA-viruses, thus, enabling heritability estimation for a broad range of pathogens.

## Introduction

Pathogens transmitted between donor and recipient hosts are genetically related much like children are related to their parents through inherited genes. This analogy between transmission and biological reproduction has inspired the use of heritability (*H*^2^) - a term borrowed from quantitative genetics (Falconer, 1996; Hartyl and Clark, 2007; Lynch and Walsh, 1998) to measure the contribution of pathogen genetic factors to pathogen traits, such as virulence, transmissibility and drug-resistance of infections.

Two families of methods enable estimating the heritability of a pathogen trait in the absence of knowledge about its genetic basis:

- Resemblance estimators measuring the relative trait-similarity within groups of transmission-related patients. Common methods of that kind are linear regression of donor-recipient pairs (DR) (Fraser *et al.*, 2014; Leventhal and Bonhoeffer, 2016) and analysis of variance (ANOVA) of patients linked by (near-)identity of carried strains (Anderson *et al.*, 2010; Shirreff *et al.*, 2013).
- Phylogenetic comparative methods measuring the association between observed trait values from patients and their (approximate) transmission tree inferred from carried pathogen sequences. Common examples of such methods are the phylogenetic mixed model (PMM) (Housworth *et al.*, 2004) and Pagel’s *λ* (Freckleton *et al.*, 2002).

Most of these methods have been applied in studies of the viral contribution to virulence of an HIV infection (Alizon *et al.*, 2010; Blanquart *et al.*, 2017; Bonhoeffer *et al.*, 2015; Fraser *et al.*, 2014; Hecht *et al.*, 2010; Hodcroft *et al.*, 2014; Hollingsworth *et al.*, 2010; Leventhal and Bonhoeffer, 2016; Lingappa *et al.*, 2013; Shirreff *et al.*, 2013; Tang *et al.*, 2004; van der Kuyl *et al.*, 2010; Yue *et al.*, 2013), quantified by log10 set point viral load – lg(spVL) – the amount of virions per blood-volume stabilizing in HIV patients at the beginning of the asymptomatic phase and best-predicting its duration (Mellors *et al.*, 1996). In the view of discrepant reports of lg(spVL)-heritability, several authors have questioned the methods’ accuracy (Fraser *et al.*, 2014; Leventhal and Bonhoeffer, 2016; Shirreff *et al.*, 2013). Shirreff et al. 2012 used simulation of trait-values on existing HIV transmission trees to reveal that phylogenetic comparative methods report strongly under- or over-estimated values depending on the true heritability value used in the simulation (Shirreff *et al.*, 2013). Later, Fraser et al. 2014 claimed that DR is unbiased with respect to lg(spVL)-heritability and is robust to trait-based selection for transmission (Fraser *et al.*, 2014). Finally, Leventhal and Bonhoeffer (2016) simulated Wright-Fisher generations of transmission confirming that DR outperforms PMM in terms of robustness and accuracy and suggesting that current phylogenetic methods are compromised by questionable assumptions, such as ultrametricity of trees (all measurements collected at the same time) and neutral evolution of the trait. These three studies assume that once the trait value is set in the recipient upon infection, it remains constant throughout its infectious time. This assumption is partially acceptable for lg(spVL), see (Geskus *et al.*, 2007) and references therein, but it is highly arguable for pathogen traits in general, because mutations during infection are often associated with phenotype changes, e.g. escape from adaptive immune response (Virgin *et al.*, 2009), drug resistance, or thermotolerance (Dessau *et al.*, 2012; Presloid *et al.*, 2016). The theory of heritability, which was developed by quantitative geneticists to study populations of animals and plants (Falconer, 1996; Hartyl and Clark, 2007; Lynch and Walsh, 1998), does not account for individual gradual evolution and other lifecycle differences between pathogens and mating species. This reveals the need for a careful transfer of the quantitative genetics terminology and methods to the domain of pathogen traits.

In the section “Overview on heritability”, we review the definitions of heritability for sexually reproducing organisms and discuss how these definitions are affected by the lifecycle differences between sexual species and pathogens. In the section “New Approaches”, we uncover the reasons for biases in current resemblance-based and phylogenetic estimators of heritability and explore two alternative approaches to overcome these biases. In the Results section, we compare the different heritability estimators using in-silico simulations of epidemics, and report a heritability analysis of spVL data from a large HIV cohort. Our results allow to establish a lower bound for the viral genetic contribution to set-point viral load. The Discussion section puts our modeling and empirical results into a broader perspective.

## Overview on heritability

### Heritability in sexual species

Jacquard (1983) noticed that the term “heritability” has been used by quantitative geneticists to serve three different concepts: (i) the genetic determination of a trait; (ii) the resemblance between relatives; (iii) the efficiency of selection. Hence, it may be confusing to use the term “heritability” without an accompanying definition or a qualifier like “narrow-sense”, “broad-sense” and “realized”. Below, we briefly introduce this terminology; formal definitions are written in the section Materials and Methods.

#### Genetic determination

Considering a real-valued (quantitative) trait, the degree to which the genes of individuals determine their trait-values is quantified in a statistical sense by the **broad-sense heritability**, *H*^2^. Assuming a sufficiently large population and full knowledge of the distinct genetic variants (genotypes) influencing the trait, *H*^2^ can be measured by the coefficient of determination, 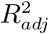, obtained over a grouping of the population by genotype. In the world of animals and plants, though, it is impossible to measure *H*^2^ in this way, because population sizes are small compared to large numbers of (usually unknown) genotypes. Thus, quantitative genetics focuses on estimating a lower bound for *H*^2^– the **narrow-sense heritability**, *h*^2^. *h*^2^ summarizes how much of the trait variance is attributable to single-locus additive genetic effects and, in sexually reproducing populations, it can be estimated from measures of the trait-resemblance between relatives.

#### Resemblance between relatives

Relatives resemble each other not only for carrying similar genes but also for living in similar environments. Hence, it is necessary to disentangle the concept of resemblance from that of genetic determination. For an ordered relationship such as parent-offspring, the resemblance is usually measured by the **regression slope**, *b*, of expected offspring values on mean parental values. For members of unordered relationships, such as identical twins, sibs and cousins, their relative resemblance is quantified by the one-way analysis of variance (ANOVA), which estimates the so-called **intraclass correlation** (ICC) denoted here as *r*_*A*_[type of relationship].

#### Efficiency of selection

The last of the three concepts is that of the efficiency of selection for breeding of the individuals with “best” trait-values. This is quantified by the **realized heritability**, 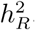, defined in Hartyl and Clark (2007) as the response to selection relative to the selection differential.

#### Connecting the dots

The success of quantitative genetics in the pre-genomic era relies on the insight that “*inferences concerning the genetic basis of quantitative traits can be extracted from phenotypic measures of the resemblance between relatives* (Lynch and Walsh, 1998)”. Mathematically, this quote is expressed as a set of approximations, which have become dogmatic in quantitative genetics:

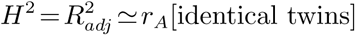

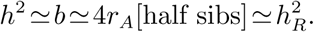

The first equation is valid in general, provided there is no strong maternal effect on the trait, the observed twins have been separated at birth and raised in independent environments and the assumptions of ANOVA such as normality and homoscedasticity are at least approximately met. The second equation, though, is provable only for diploid sexually reproducing species. This is because genetic segregation and recombination during sexual reproduction ensure that single-locus additive effects are inherited at bigger proportions (1/2 from each parent) compared to multi-locus (epistatic) interactions (i.e. 1/4 for 2-loci-, 1/8 for 3-loci-interactions, etc) (Falconer, 1996; Lynch and Walsh, 1998).

In summary, in sexually reproducing populations, heritability is used to quantify to what extent the genetics explain a trait (broad-sense heritability, *H*^2^) as well as to measure or predict the response to trait-based selection for reproduction (realized heritability, 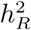). Since it is practically hard to measure *H*^2^, one often uses empirical measures of the resemblance between relatives (i.e. parent-offspring regression, *b*, or ICC from half sibs, *r*_*A*_) to estimate the extent, to which single-locus additive effects determine the trait (narrow-sense heritability, *h*^2^). It turns out that 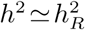, justifying the dual role of *h*^2^ as a measure of genetic determination and a measure for the rate of trait-evolution resulting from selection.

Transfer to pathogen traits

The transfer of the above terminology from traits of diploid organisms to pathogen traits is almost verbatim and only requires substituting “pathogen genes” for “organism genes”, “donor value” for “parental value” and “recipient value” for “offspring value”. However, three important differences between the lifecycles of diploid organisms and pathogens alter the connections between the definitions and the estimators:

- **Asexual haploid nature of pathogen transmission** The first difference is that, unlike reproduction of diploid organisms, the transmission of a pathogen from a donor to a recipient is more similar to asexual reproduction in haploid organisms, because, typically, whole pathogens get transferred between hosts. Importantly, in the absence of genetic segregation and recombination at transmission, there is no preference in transmitting single-locus over multi-locus genetic effects.
- **Partial quasispecies transmission** The second difference is that the transmitted proportion of genetic information characterizing the pathogen in the donor is unknown and varying between transmission events. For example, for slowly evolving bacteria such as *Micobacterium tubercolosis* (Mtb), transmission can be clonal (Bjorn-Mortensen *et al.*, 2016), whereas, for rapidly evolving retroviruses like HIV, transmission is often accompanied by bottlenecks causing only a tiny sample of the large and genetically diverse virus population in the donor (aka quasispecies) to penetrate and survive in the recipient (Keele *et al.*, 2008).
- **Within-host pathogen evolution** The third difference involves the change in phenotypic value due to within-host pathogen mutation and recombination. While genetic change is rare during the lifetime of animals and plants and its phenotypic effects are typically delayed to the offspring generations, it constitutes a hallmark in the lifecycle of pathogens and causes a gradual or immediate phenotypic change such as increasing virulence, immune escape or drug resistance.

For equal genotypes in donor and recipient as well as for distributions of donors and recipients being equal to the total population distribution, the estimators *b* and *r*_*A*_ evaluated on transmission pairs would be unbiased with respect to *H*^2^. This has been shown in theory (Fraser *et al.*, 2014). Further, Leventhal and Bonhoeffer (2016) showed through simulations that DR is accurate in the case of minute evolution in the recipient host upon infection. In their simulation, partial quasispecies transmission and gradual within-host evolution throughout the infection is ignored. We notice, though, that these two phenomena cause a negative bias in *b* and *r*_*A*_ as estimator of *H*^2^, because they co-act for the loss of resemblance without affecting *H*^2^ in any way. Thus, *b* and *r*_*A*_ should be regarded as statistics summarizing the resemblance in transmission couples observable after partial quasispecies transmission and delay between transmission and measurements. Further in the text, we use the symbols *b*_*τ*_ and *r*_*A,τ*_ to emphasize that these estimators have been calculated on a sample of donors and recipients with (variable) periods *τ*_*d*_ and *τ*_*r*_ between transmission and measurements, *τ* = *τ*_*d*_ +*τ*_*r*_ denoting the total amount of time between measurements (fig. 1). By contrast, we use *b*_0_ and *r*_*A,*0_ to emphasize that the calculation has been done on the immediate trait-values right after transmission.

**FIG. 1.**
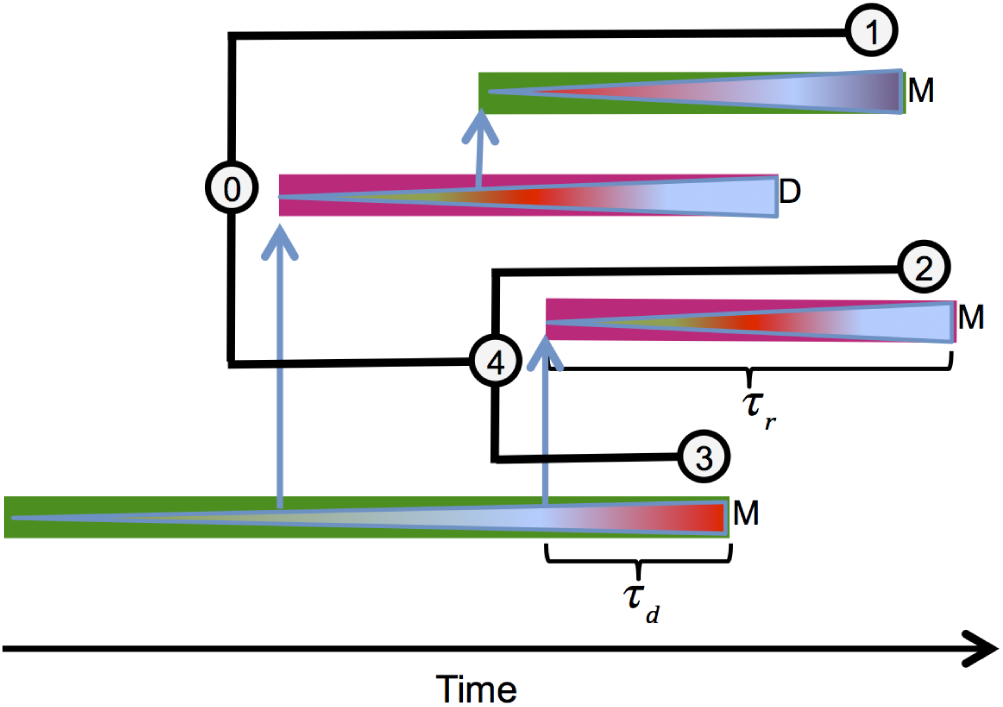
A schematic representation of an epidemic. Colored rectangles represent infectious periods of hosts, different colors corresponding to different host types. Triangles inside hosts represent pathogen quasispecies, change of color indicating substitution of dominant strains. Capital letters denote host-events: M: phenotype measurement and recovery; D: host death. Vertical arrows show the time and direction of transmission events. The within-donor and within-recipient measurement delays, *τ*_*d*_ and *τ*_*r*_, are shown for one donor-recipient couple. The transmission tree connecting the measured hosts is drawn in black (shifted up-left for visualization purpose). Notice that, due to incomplete sampling, some infectious periods (colored rectangles) are shorter than their corresponding lineages on the tree. Couples of tips that are each other’s closest tip by phylogenetic distance, e.g. (2,3), are called “phylogenetic pairs” (PPs). With some exceptions, PPs coincide with pairs of tips descending from the same parent node (a.k.a. siblings or “cherries”).

#### Phylogenetic heritability

As an alternative to resemblance-based methods, it is possible to fit a parametric model of the trait-evolution along the branches of the transmission tree connecting the patients (fig. 1). For example, the phylogenetic mixed model (PMM) (Housworth *et al.*, 2004; Lynch, 1991) assumes an additive model of the trait-values, *z*(*t*) = *g*(*t*)+*e*, in which *z*(*t*) represents the trait-value at time *t* for a given lineage of the tree, *g*(*t*) represents a heritable (genotypic) value at time *t* for this lineage and *e* represents a non-heritable contribution representing the sum of cumulative environmental effects on the trait and measurement error.

The PMM assumes that *g*(*t*) evolves along the tree according to a branching Brownian motion process defined by the stochastic differential equation:

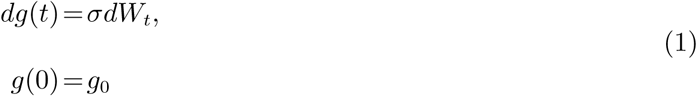

where *g*_0_ is the initial genotypic value at the root, *W*_*t*_ is the standard Wiener process and *σ >* 0 is the unit-time standard deviation (Grimmett and Stirzaker, 2001).

The environmental contribution *e* can change along the tree in any way as long as the values *e* at the tips are independent and identically distributed (i.i.d.) normal with mean 0 and variance 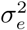. In the case of an epidemic, *e* represents the contribution from an individual’s immune system; it obtains a value at the beginning of an infection, which can stay constant or change during the course of an infection, but is uncorrelated to the immune systems of other hosts. Denoting by 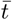 the mean root-tip distance in the tree, the **phylogenetic heritability** is defined as the expected proportion of phenotypic variance attributable to *g* at the tips:

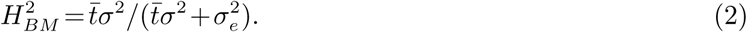

For rapidly evolving pathogens, such as RNA viruses, it is possible to infer the approximate transmission tree from pathogen sequences sampled at the moment of trait measurement (Hu *et al.*, 2004). This has inspired the use of PMM to estimate lg(spVL)-heritability in HIV patients (Alizon *et al.*, 2010; Hodcroft *et al.*, 2014; Shirreff *et al.*, 2013). However, this approach has been questioned in recent simulation tests reporting strongly positively or negatively biased PMM estimates with respect to the simulated *H*^2^ (Leventhal and Bonhoeffer, 2016; Shirreff *et al.*, 2013).

#### Summary

In summary, for pathogen traits, measures of resemblance, such as *b*_0_ and *r*_*A,*0_, should be considered as estimates of *H*^2^, compromised by quasispecies differences, rather than estimates of *h*^2^. In the absence of genetic segregation and recombination at transmission, *h*^2^ loses its dual role as an accessible measure of genetic determination and as a predictor for the rate of evolution. Due to delayed diagnosis, data from transmission couples for estimating *b*_0_ and *r*_*A,*0_ is rarely available in practice, while *b*_*τ*_ and *r*_*A,τ*_ are negatively biased due to gradual within-host evolution. Phylogenetic methods, such as PMM, should provide an alternative for estimating *H*^2^ but recent simulation tests suggest that these methods are not well suited to the study of pathogen traits.

### New Approaches

In this section, we first show through a real world example that the current methods, both resemblance-based and phylogenetic, are prone to strong negative bias in estimating *H*^2^. As a principal cause, we reveal the inability of these methods to model the gradual loss of phenotypic resemblance between transmission related patients as a function of their phylogenetic distance. Then, we propose two alternative approaches to account for this phenomenon. Finally, we design a toy model of an epidemic, which we use as a validation tool for the different heritability estimators.

#### Uncovering biases in current heritability estimators

Previous studies of malaria and HIV have used clustering of the tips in the transmission tree to identify donor-recipient couples (Hecht *et al.*, 2010; Hollingsworth *et al.*, 2010; Shirreff *et al.*, 2013) or groups of transmission related patients (Anderson *et al.*, 2010). In particular, Shirreff *et al.* (2013) defines the method of phylogenetic pairs (PP) as ANOVA on pairs of tips in the transmission tree that are mutually nearest to each other by phylogenetic distance (*τ*) (fig. 1). Taking this approach a step further, we order the PPs by *τ* and split them into bins of equal size, evaluating the correlation between pair trait-values (*r*_*A*_) in each bin. An analysis of 1912 PPs extracted from a recently published transmission tree of 8473 HIV patients (Hodcroft *et al.*, 2014) reveals a well pronounced decrease of the correlation between pair-values (black points and vertical bars on fig. 2). For small *τ* (left-most bin), the correlation *r*_*A*_ is far above the 95% CI estimated by the PP-method (thick grey horizontal bar), while, for big values of *τ*, *r*_*A*_ falls below the 95% CI estimated by the PP-method. A similar pattern is observed when applying DR (using *b* instead of *r*_*A*_ upon assigning a donor and a recipient at random in each phylogenetic pair; results not shown). Being ignorant of *τ*, all resemblance-based methods average over *τ* in the observed sample of pairs. Thus, these methods should be considered negatively biased in general. They can approximate the true *H*^2^ in the population only in the limit *τ →* 0 and up to additional sources of bias such as partial quasispecies transmission and differences in the distributions of donors, recipients and total population. Further, we repeatedly simulate trait-values on the transmission tree under the maximum likelihood fit of the PMM method and re-evaluate the correlation in the same bins of PPs. Plotting the resulting correlation estimates from the simulations next to the correlation in the original data shows that PMM does not reproduce the gradual loss of correlation as a function of *τ* (brown points and vertical bars on fig. 2). To understand the reason for that, we consider the initial assumption of the PMM method. According to Brownian motion, the covariance between the values of a pair of tips (*ij*) is proportional to the distance *t*_*ij*_ from the root to their most recent common ancestor (mrca):

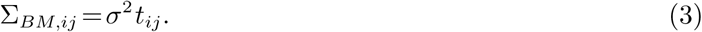

Without an additional requirement for ultrametricity of the tree (all tips at equal distance from the root), this assumption does not imply a relationship between the covariance and the phylogenetic distance between the tips. In real non-ultrametric transmission trees, though, we observe a rapid loss of covariance as *τ* increases, while there is only a weak relationship between covariance and root-mrca distance. The latter is reflected also by a nearly horizontal slope of the expected covariance between PPs at distance *τ* modeled under PMM as a function of their mean root-mrca distance 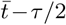,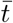 denoting the mean root-tip distance (brown line on fig. 2). We conclude that the BM assumption is inappropriate for modeling the evolution of pathogen traits along transmission trees. Instead, we need to model the covariance between trait-values at the tips as a function of their phylogenetic distance.

**FIG. 2.**
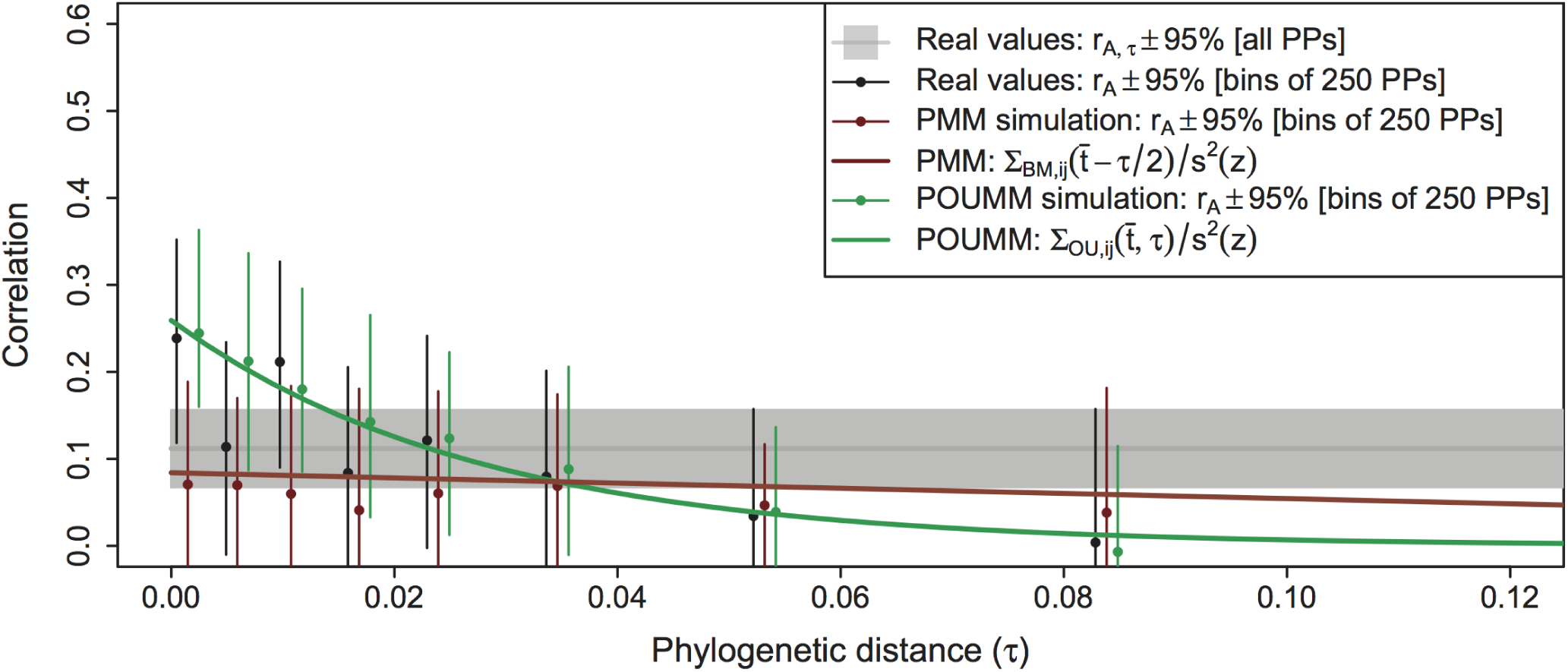
Gradual loss of phenotypic resemblance between transmission related patients. A sample of PPs with real lg(spVL)-measurements from HIV patients shows a decrease in the correlation (ICC) between pair trait-values as a function of the pair phylogenetic distance *τ*. The PPs have been ordered according to *τ* and split into bins of 250 PPs (i.e. 500 patients). A thin and a thick grey horizontal bars denote the correlation with 95% CI in all PPs; black points with vertical bars denote the correlation with 95% CI within each bin. Brown and green points with vertical bars denote the mean correlation and 95% CI obtained after replacing the real trait values on the tree by values simulated under the maximum likelihood fit of the PMM and the POUMM methods respectively (mean and 95% CI estimated from 40 replications). A brown and a green line show the expected correlation between pairs of tips at distance *τ* as modeled by the PMM and the POUMM method: for PMM, this is the normalized covariance ∑_*BM,ij*_ at the mean distance 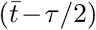*-τ /*2 from the root to the pair’s most recent common ancestor (eq. 3); for POUMM, this is the normalized covariance ∑_*OU,ij*_ at the mean root-tip distance 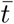 and tip-pair distance *τ* (eq. 7). To approximate correlation, both covariances have been normalized by the total phenotypic variance, *s*^2^(*z*).

#### New estimators of pathogen trait heritability

*ANOVA on closest PPs.* Assuming that the correlation measured in the bin at minimal *τ* is a more accurate approximation of *H*^2^ than the correlation in bins at bigger *τ* or the correlation of all PPs, we refine the PP-method by imposing a limit on *τ* and define closest phylogenetic pairs (CPP) as PPs that are not farther apart than a cut-off distance *τ*^′^. We tune this parameter based on the trade-off that arises between the negative bias caused by *τ* and the loss of statistical power caused by omitting data. Further in the text, we refer to this variant of the PP method as **ANOVA-CPP** with estimate *r*_*A,τ*′_. The main drawback of this filtering technique is its reduced statistical power due to fewer observations.

*The phylogenetic Ornstein-Uhlenbeck mixed model.* The phylogenetic Ornstein-Uhlenbeck mixed model (POUMM) is an extension of the PMM replacing the BM assumption with an assumption of an Ornstein-Uhlenbeck (OU) process for the genotype evolution (Mitov and Stadler, 2017). The OU-process represents a continuous time random walk, which tends to move around a long-term mean value with greater attraction when the process is further away from that value (Uhlenbeck and Ornstein, 1930). Technically, this is accomplished by adding an attraction term to eq. 1:

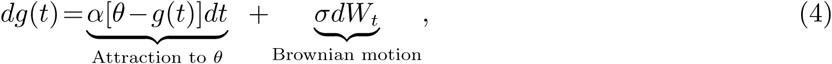

where *θ* denotes the long-term mean and *α >* 0 is the attraction strength. Since in the limit *α →* 0 the attraction term vanishes and only the BM term remains, the OU-process represents a generalization of BM. As in the PMM, a white noise is added to *g*(*t*) at the tips. POUMM estimates the parameters of the stochastic model and the white noise, and then evaluates the phylogenetic heritability as a function of 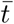:

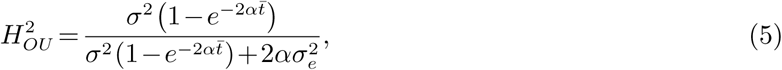

or as a time-independent function of the trait’s sample variance, *s*^2^(*z*) (Mitov and Stadler, 2017):

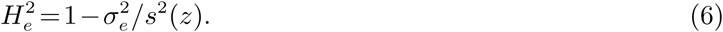

Further in the text, we use the symbols 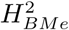 and 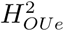 to denote the time-independent heritability (eq.6) inferred respectively by PMM and POUMM.

The POUMM provides an interesting alternative to the PMM, since, under this model, the expectation for the covariance between trait-values for a couple of tips (*ij*) is a function of both, their root-mrca distance, *t*_*ij*_, and their phylogenetic distance *τ*_*ij*_:

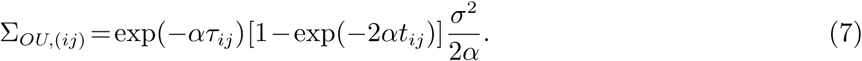

As it turns out, data simulated under the maximum likelihood fit of the POUMM method reproduces the loss of resemblance between PPs in the UK HIV data (green points and vertical bars on fig. 2). Plotting the correlation between tips in the tree as expected by the OU-process (substituting 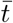 for *t*_*ij*_ in eq. 7 and normalizing by *s*^2^(*z*); green line on fig. 2) reveals that the correlation between transmission-related patients decreases approximately exponentially with rate-constant equal to to the parameter *α* of the OU process, ML estimate *α* = 36.3, 95% CI [22.6, 62.4].

#### A toy-model of an epidemic

To test different estimators of heritability, we implement a phenomenological model of an epidemic, in which an imaginary pathogen trait, *z*, is determined by the interaction between the alleles at a finite number of loci in the pathogen genotype and a finite number of immune system types encountered in the susceptible population. This toy-model is embedded into a stochastic Susceptible-Infected-Recovered (SIR) epidemic model (Keeling and Rohani, 2007), implementing “neutral” and “selection” modes of within- and between-host dynamics. With the aid of figure 3, we briefly describe this model, leaving the technical details for the section Materials and Methods.

**FIG. 3.**
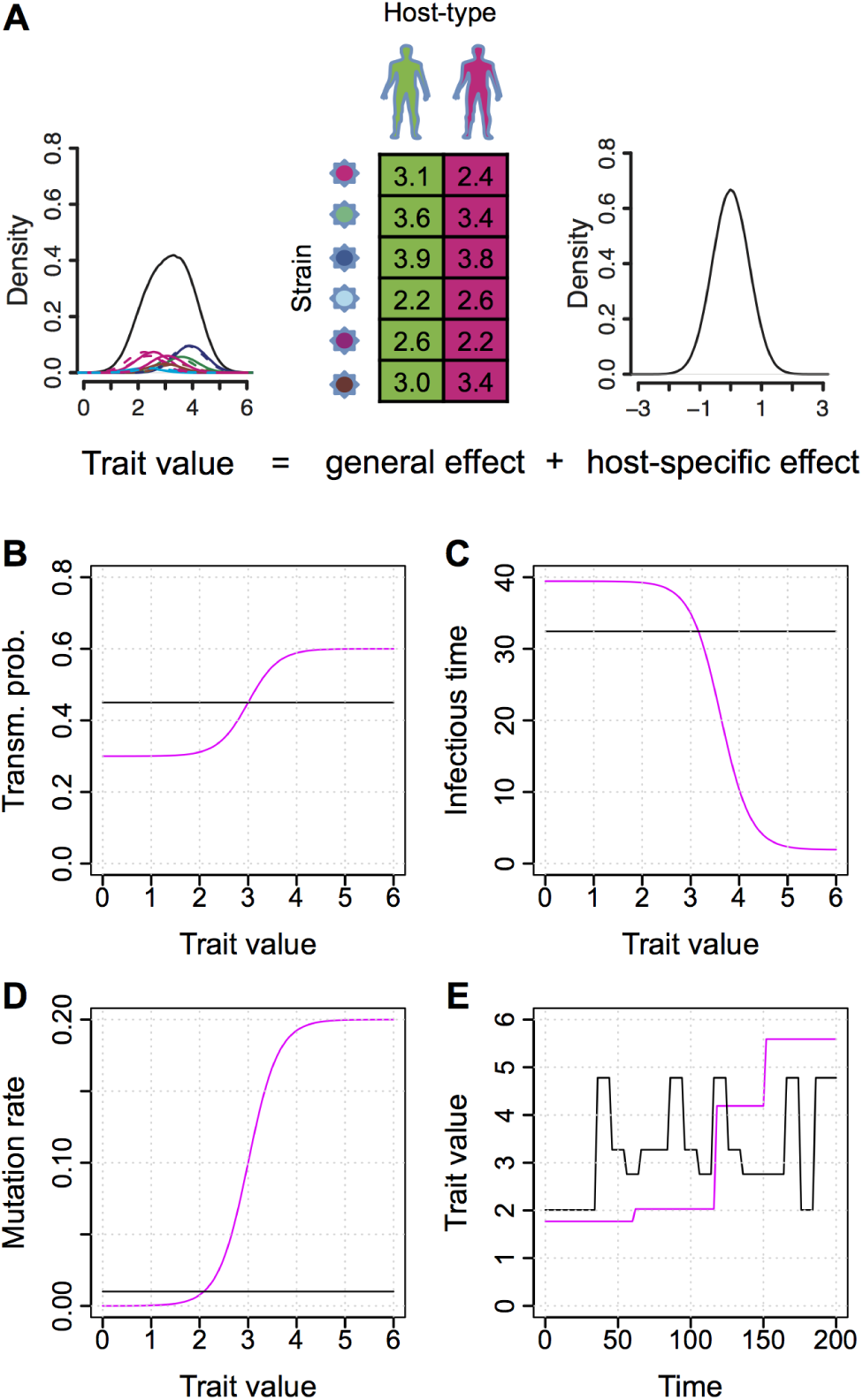
A toy model of a pathogen trait with within-host evolution and SIR dynamics. (A) Schematic representation of a pathogen trait formed from a general ¡host type *×*carried strain¿ effect and a host-specific effect. The density of the trait-values in an infected population represents a mixture of normal densities corresponding to each one of twelve host type*×* strain combinations, scaled by their frequencies (dashed-lines depict host type 2); (B-E) SIR dynamics, color indicating selection modes with respect to to the trait-value, black: neutral, magenta: select (as specified in table 1); (B) Per risky contact transmission probability; (C) Expected infectious period if no mutation happens; (D) mutation rate (in mutations per site per time unit); (E) Example time-course of the trait evolution within a host; Time and trait-value units are arbitrary.

We assume two equally frequent and lifelong immutable types of host immune system and two mutable trait-determining loci in the pathogen genotype. With *M*_1_ = 3 and *M*_2_ = 2 possible alleles at each locus, there are six possible genotypes denoted 1:11, 2:12, 3:21, 4:22, 5:31, 6:32 (fig. 3A). We assume absence of strain coexistence within a host, so that the within-host quasispecies is represented by a single strain. At a time *t*, the value *z*_*i*_(*t*) of an infected individual *i* is defined as a function of its immune system type, **y**_*i*_ ∈ {1,2}, the currently carried strain **x**_*i*_(*t*) ∈ {1,…,6}, and the individual’s specific effect for this strain *e*_*i*_[**x**_*i*_(*t*)] ∼ 𝒩 (0, 0.36) drawn at random for each strain (in each infected individual). We call a (type **y**-**x**) **general effect** the expected trait value of type-**y** carriers of strain **x** in an infected population: *GE*[**y**,**x**] = E[*z|***y**,**x**]. For a set of fixed general effects, *z*_*i*_(*t*) is constructed according to the equation:

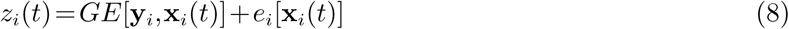

**Table 1.** Within- and between-host dynamics of the toy epidemiological model.

| Scope | Parameter | neutral | select |
| --- | --- | --- | --- |
| Between-host | Natural birth rate | $\lambda_{\text{nat}} = 117.6$ | |
| | Natural per capita death rate | $\delta_{\text{nat}} = 1/850$ | |
| | Per capita recovery rate | $\rho = 1/48$ | |
| | Per capita contact rate | $\kappa \in \{\frac{1}{2}, \frac{1}{4}, \frac{1}{6}, \frac{1}{8}, \frac{1}{10}, \frac{1}{12}\}$ | |
| | Per capita risky contact rate (S: current proportion of susceptible in the pop.) | $S \times \kappa$ | |
| | Per risky contact transmission probability | $\gamma_{\text{neutral}} = .45$ | $\gamma(z) = \gamma_{\min} + \frac{(\gamma_{\max} - \gamma_{\min})(\gamma_{50})^{\gamma_k}}{10^{z\gamma_k} + (\gamma_{50})^{\gamma_k}}$, where $\gamma_{\min} = .3, \gamma_{\max} = .6, \gamma_{50} = 10^3, \gamma_k = 1.4$ |
| | Per capita death rate for infected individuals | $\delta_{\text{neutral}} = .01$ | $\delta(z) = \delta_{\text{nat}} + \frac{10^{zD_k} + (D_{50})^{D_k}}{D_{\min} 10^{zD_k} + D_{\max} (D_{50})^{D_k}}$, where $D_{\min} = 2, D_{\max} = 300, D_{50} = 10^3, D_k = 1.4$ |
| Within-host | Per locus pathogen mutation rate | $\nu_{\text{neutral}} = .01$ | $\nu(z) = \frac{\nu_{\max}(\nu_{50})10^{z\nu_k}}{10^{z\nu_k} + (\nu_{50})^{\nu_k}}$, where $\nu_{\max} = .2, \nu_{50} = 10^3, \nu_k = 1.4$ |
| | Rate of substitution of strain $\mathbf{x}_j$ for $\mathbf{x}_i$, where $\mathbf{x}_i \neq \mathbf{x}_j$ at a single locus, $l$, $M_l$ is the number of alleles at locus $l$, and the corresponding values are $z_i$ and $z_j$ | $\xi_l = \frac{\nu_{\text{neutral}}}{M_l - 1}$ | $\xi_{l,i \leftarrow j}(z_i, z_j) = \begin{cases} \frac{\nu(z_i)}{M_l - 1} & \text{if } \nu(z_i) < \nu(z_j) \\ 0 & \text{, otherwise} \end{cases}$ |

We use a fixed set of general effects drawn from the uniform distribution *U* (2,4) for the twelve **y**-**x** combinations (fig. 3A).

We embed this trait-model into a stochastic Susceptible-Infected-Recovered (SIR) model of an epidemic with demography and frequency dependent transmission as described in (Keeling and Rohani, 2007), ch. 1. Each infected individual, *i*, has a variable trait value *z*_*i*_(*t*) constructed as in eq. 8. Within-host phenomena (strain mutation and substitution) and between-host phenomena (natural birth, contact, transmission, diagnosis, recovery and death) occur at random according to Poisson processes. The rate parameters defining these processes are written in table 1.

For each group of parameters (within- and between-host), we consider the following two modes of dynamics:

- neutral: rates are defined as global constants mimicking neutrality (i.e. lack of selection) with respect to *z* (black lines on fig. 3B-D). For within-host phenomena, it is assumed that a mutation of the pathogen is followed by instantaneous substitution of the mutant for the current dominant strain, regardless of the induced change in *z* (black line on fig. 3E);
- select: borrowing the approach from (Fraser *et al.*, 2007), the rates of transmission and within-host pathogen mutation are defined as increasing Hill functions of 10^*z*^, while the infected death rate is defined as an inverse decreasing Hill function of 10^*z*^, thus mimicking increasing per capita transmission- and pathogen-induced mortality for higher *z* (red lines on fig. 3B-D). Within hosts, it is assumed that a mutation of the pathogen is followed by instantaneous substitution only if it resulted in a higher *z*. Otherwise, the mutation is considered deleterious (red line on fig. 3E).

By combining “neutral” and “select” dynamics for the strain mutation and substitution rates at the within-host level, and the virus-induced per capita death rate and per contact transmission probability at the between-host level, we define the following four scenarios (fig. 4):

- Within: neutral / Between: neutral;
- Within: select / Between: neutral;
- Within: neutral / Between: select;
- Within: select / Between: select;

**FIG. 4.**
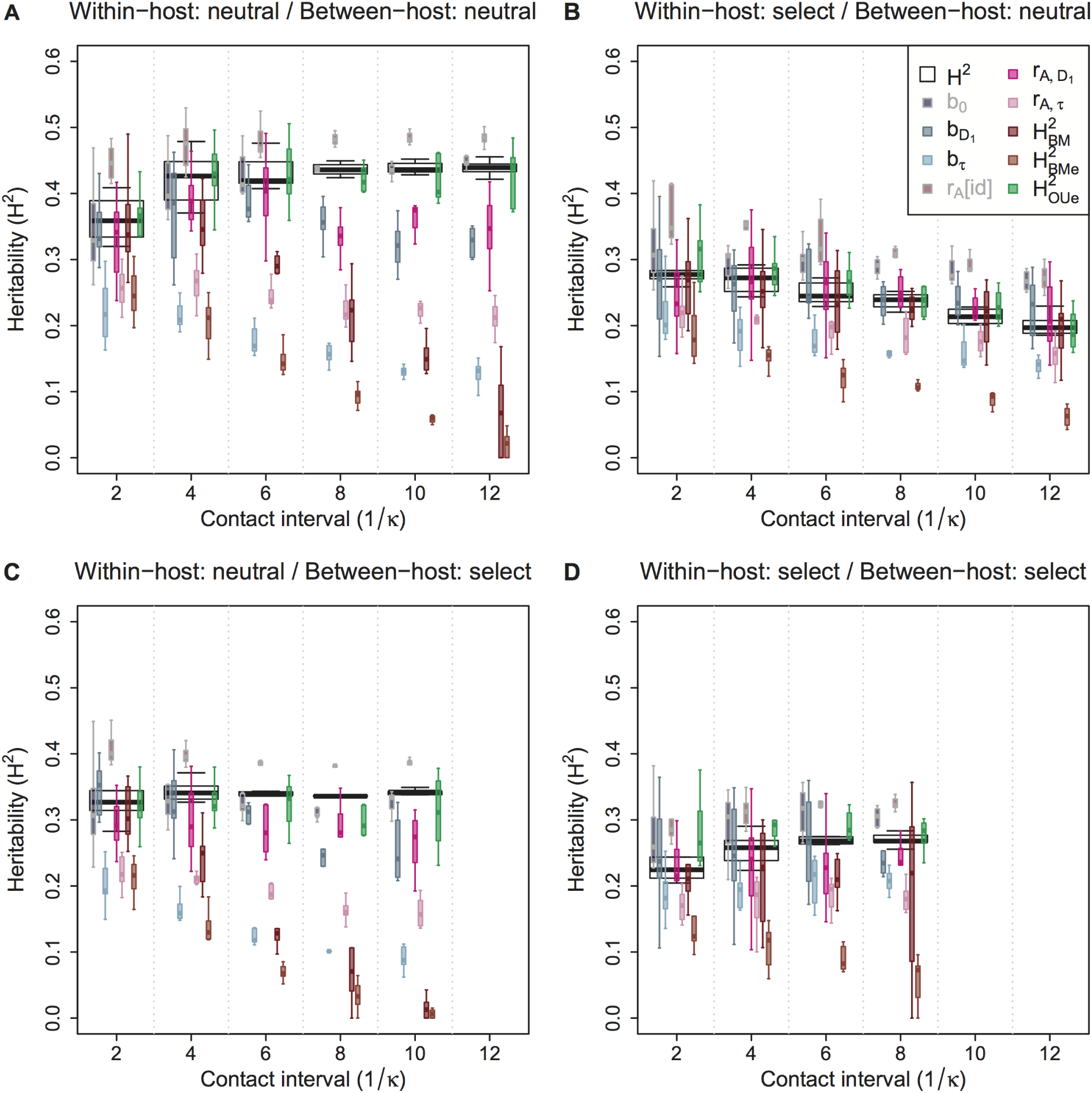
Heritability estimates from computer simulations of the toy-model. (A-D) *H*^2^-estimates in simulations of “neutral” and “select” within-/between-host dynamics. Each box-group summarizes simulations (first up to 10,000 diagnoses) at a fixed contact rate, *κ*; white boxes (background) denote true heritability, colored boxes denote estimates (foreground). Estimates that are not measurable in practice are shaded in grey. Statistical significance is evaluated through t-tests summarized in table 2. For an in-depth analysis of bias for fixed contact-rate *κ*, see Supplementary Notes, supplementary figs. S1 and S2. Estimates of 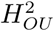 are not shown since they match with 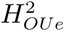.

**Table 2.** Mean difference 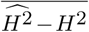 from the toy-model simulations grouped by scenario. Statistical significance is estimated by Student’s t-tests, p-values denoted by an asterisk as follows: * *p <* 0.01; ** *p <* 0.001. The column # denotes the number of simulations resulting in an epidemic outbreak under each scenario. Grey background indicates estimates that are unavailable in practice. Estimates of 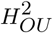 are not shown since they match with 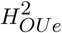.

| Within/Between | # | $b_0$ | $b_{D_1}$ | $b_\tau$ | $r_A[id]$ | $r_{A,D_1}$ | $r_{A,\tau}$ | $H_{BM}^2$ | $H_{BMe}^2$ | $H_{OUe}^2$ |
| --- | --- | --- | --- | --- | --- | --- | --- | --- | --- | --- |
| neutral/neutral | 50 | -0.01* | -0.07** | -0.25** | 0.05** | -0.05** | -0.18** | -0.17** | -0.28** | -0.01 |
| select/neutral | 47 | 0.05** | 0 | -0.07** | 0.08** | 0 | -0.06** | -0.01 | -0.12** | 0.01* |
| neutral/select | 41 | -0.02** | -0.04** | -0.2** | 0.05** | -0.06** | -0.15** | -0.17** | -0.24** | -0.02** |
| select/select | 37 | 0.04** | -0.01 | -0.06** | 0.06** | -0.03* | -0.08** | -0.04* | -0.16** | 0.03** |

For each of these scenarios and mean contact interval 1*/κ ∈ {*2, 4, 6, 8, 10, 12*}* (arbitrary time units), we perform ten simulations resulting in a total of 4*×*6*×*10 = 240 simulations. In the next section, we discuss the resulting heritability estimates from these simulations and from a real dataset.

## Results

### Simulations

Of 240 toy-model simulations, 175 resulted in epidemic outbreaks of at least 1,000 diagnosed individuals. In each of these 175 simulations, we analyzed the population of the first up to 10,000 diagnosed individuals. We denote this population by *Z*_10k_ and the corresponding transmission tree – by *T*_10k_.

The direct measure of broad-sense heritability, 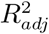, was compared to the following estimators: *b*_0_ in all transmission couples found in *Z*_10k_; *b*_*τ*_ in the same transmission couples; *b*_*D*_1 in transmission couples in *Z*_10k_ having *τ* not exceeding the first decile, *D*_1_; *r*_*A*_[id] based on grouping by identity of carried strain in *Z*_10k_; *r*_*A,τ*_ based on phylogenetic pairs (PPs) in *T*_10k_; *r*_*A,D*1_ based on closest phylogenetic pairs (CPPs) defined as PPs in *T*_10k_ having *τ* not exceeding the first decile, *D*_1_, among all PPs; 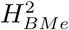 and 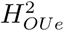 based on the maximum likelihood (ML) fit of the PMM and POUMM methods on *T*_10k_. To calculate *b*_0_, we used the immediate trait-values at moments of transmission (usually not available in practice). All other estimators were calculated using trait-values at the moment of diagnosis.

A detailed analysis of the different heritability estimates (table 2, fig. 4, Supplementary Notes, supplementary figs. S1, S2, S3) confirmed the negative bias due to measurement delays in the resemblance-based estimators *b*_*τ*_ and *r*_*A,τ*_. This bias was increasing with the mean contact interval, 1*/κ*, because, for a fixed recovery rate *ρ*, rarer transmission events resulted in longer transmission trees and, therefore, longer average phylogenetic distance between tips, *τ*, (fig. S3). The negative bias was far less pronounced when imposing a threshold on *τ*, but this came at the cost of statistical power (more accurate but longer box-whisker plots for *b*_*D*_1__ and *r*_*A,D*_1__ compared to *b*_*τ*_ and *r*_*A,τ*_, fig. 4). Further, the simulations showed that a worsening fit of the BM model on longer transmission trees caused an inflated estimate of the environmental variance, 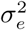, in the PMM method and, therefore, a negative bias in 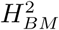 and 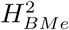. As explained in the previous section, this is caused by the inability of the BM assumption to model the loss of phenotypic resemblance with increasing phylogenetic distance between tips. Several other sources of bias, such as non-linear dependence of recipient on donor-values and deviation from normality were identified and are summarized in table 3. We conclude that, apart from the practically inaccessible immediate donor-recipient regression (*b*_0_) and ICC of patients grouped by identity of carried strain (*r*_*A*_[id]), the most accurate estimator of *H*^2^ in the toy-model simulations is 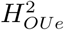 followed by estimators minimizing measurement delays such as *b*_*D*_1__ and *r*_*A,D*_1__.

**Table 3.** Sources of bias in estimators of *H*^2^. The bias direction is indicated by a “+” or a “–”, separated by a “/” when both directions are possible. The number of signs indicates the relative intensity of the bias that is observed in the simulations or in the analysis of the HIV data. A zero indicates no bias observed. A “?” indicates unknown direction. Horizontal lines separate sources that were identified in the SIR simulations (top) from sources identified in the analysis of the HIV data (middle) and sources suggested by this or previous works that were not tested (bottom). Grey background denotes estimators not available in practice.

| Source of bias | $b_0$ | $b_{D_1}$ | $b_\tau$ | $r_A[id]$ | $r_{A,D_1}$ | $r_{A,\tau}$ | $H_{BM}^2$ | $H_{BMe}^2$ | $H_{OU}^2$ | $H_{OUe}^2$ |
| --- | --- | --- | --- | --- | --- | --- | --- | --- | --- | --- |
| Gradual within-host evolution | 0 | − | − − | 0 | − | − − | 0 | 0 | 0 | 0 |
| Violation of assumed model (i.e. BM) | 0 | 0 | 0 | 0 | 0 | 0 | +/− − | +/− − | +/− | +/− |
| Nonlinear dependence of expected recipient value on donor value | +/− | +/− | +/− | 0 | 0 | 0 | 0 | 0 | 0 | 0 |
| Non-normality of $z$ | +/− | +/− | +/− | +/− | +/− | +/− | +/− | +/− | +/− | +/− |
| Loss of phylogenetic signal due to scarce transmission tree | 0 | 0 | 0 | 0 | 0 | 0 | +/− | +/− | +/− | +/− |
| Outliers in $z$ | +/− | +/− | +/− | +/− | +/− | +/− | +/− | +/− | +/− | +/− |
| Partial quasi-species transmission | − | − | − | 0 | − | − | − | − | − | − |
| Temporal changes in the trait distribution | +/− | +/− | +/− | +/− | +/− | +/− | +/− | +/− | +/− | +/− |
| Outlying tips in the tree | 0 | 0 | 0 | +/− | +/− | +/− | +/− | +/− | +/− | +/− |
| Non-homogeneous evolutionary process | 0 | 0 | 0 | 0 | − | − | +/− | +/− | +/− | +/− |
| Random error in transmission tree | 0 | 0 | 0 | 0 | 0 | 0 | − | − | − | − |
| Biases in the transmission tree | 0 | 0 | 0 | 0 | ? | ? | ? | ? | ? | ? |

#### Analysis of HIV-data

We performed ANOVA-CPP and POUMM on data from the UK HIV cohort comprising lg(spVL) measurements and a tree of viral (pol) sequences from 8,483 patients inferred previously in (Hodcroft *et al.*, 2014). The goal was to test our conclusions on a real dataset and compare the *H*^2^-estimates from ANOVA-CPP and POUMM to previous PMM/ReML-estimates on exactly the same data (Hodcroft *et al.*, 2014). A scatter plot of the phylogenetic distances of tip-pairs against the absolute phenotypic differences, *|*Δ lg(spVL)*|*, reveals a small set of 116 PPs having *τ* ⩽ 10^−4^ while the phylogenetic distance in all remaining tip-pairs is more than an order of magnitude longer, i.e. *τ >*10^−3^ (fig. 5A). A box-plot graph of the trait-values along the tree shows that the range of trait-values is confined between 1 and 7 with relatively stable median and interquartile range (IQR) throughout the epidemic (fig. 5B). This visual analysis of the data suggests that the distribution of trait values has been at equilibrium during the time period covered by the transmission tree. The random distribution of the CPPs along the transmission tree suggests that these phylogenetic pairs correspond to randomly occurring early detections of infection (trait-values from each pair depicted as magenta segments on fig. 5B). Based on the observed gap of *τ*, we defined these PPs as closest ones (CPP). We applied the 1.5*×IQR*-rule on *|*Δ lg(spVL)*|* to identify outliers among the CPPs. According to this rule, outliers are all CPPs having absolute phenotypic difference below *Q*_1_ *−*1.5*×IQR* or above *Q*_3_ +1.5*×IQR*, *Q*_1_, where *Q*_3_ denotes the 25^th^ and 75^th^ quantile of *|*Δ lg(spVL)*|* in CPPs and IQR denotes the interquartile range Q3-Q1. The outlier CPPs defined in that way are shown as blue bullets on fig. 5.

**FIG. 5.**
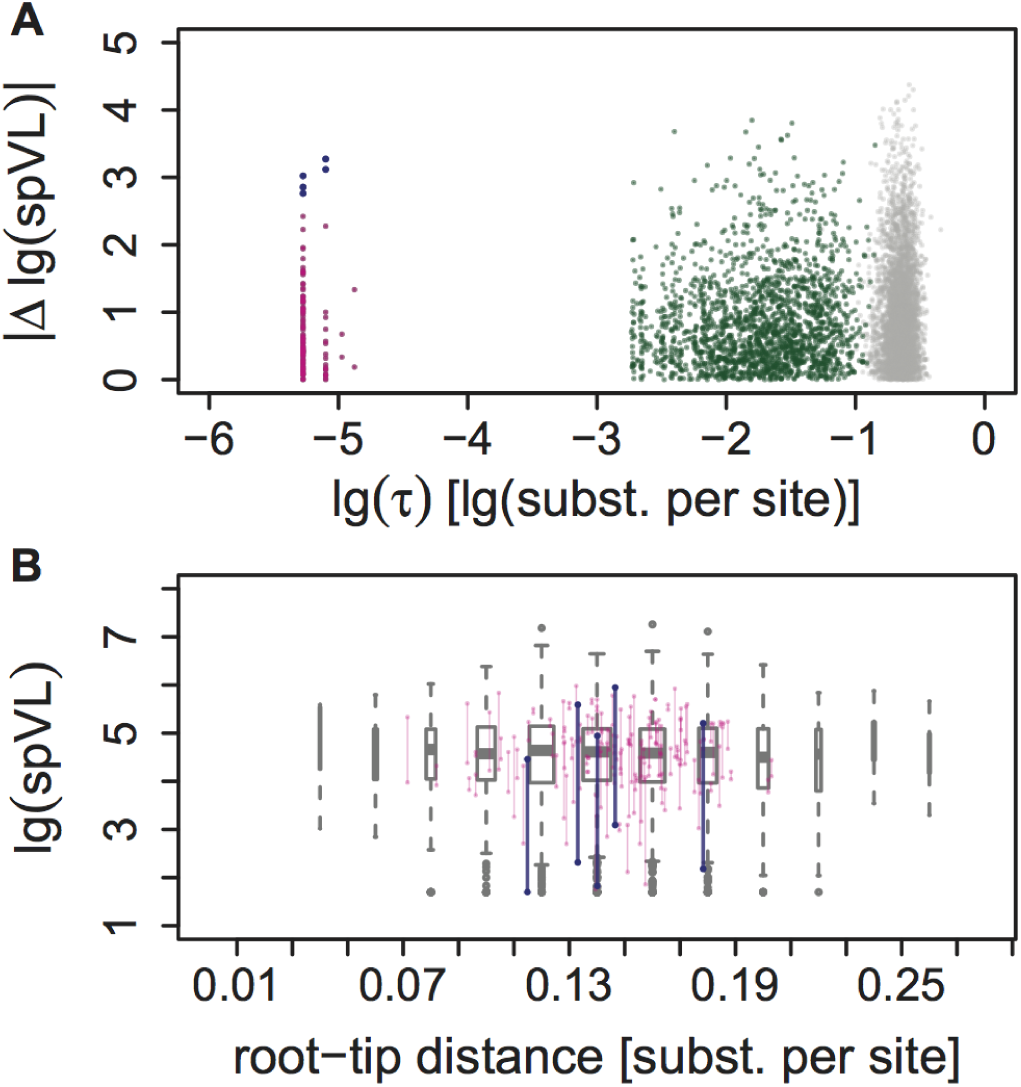
Analysis of HIV-data from UK. (A) A scatter plot of the phylogenetic distances between pairs of tips against their absolute phenotypic differences: grey – random pairs; green – PPs (*τ >*10^−4^); magenta: CPPs (*τ* ⩽ 10^−4^); blue – outlier CPPs (CPPs, for which *|*Δ lg(spVL) *|>Q*3 +1.5*×* (*Q*3 *−Q*1), *Q*1 and *Q*3 denoting the 25^th^ and 75^th^ quantile of *|*Δ lg(spVL) *|* among CPPs); (B) A box-plot representing the trait-distribution along the transmission tree. Each box-whisker represents the lg(spVL)-distribution of patients grouped by their distance from the root of the tree measured in substitutions per site. Wider boxes indicate groups bigger in size. Bullet-ending segments denote lg(spVL)-values in CPPs.

We compared the following estimators of *H*^2^, with and without inclusion of outlier CPPs in the data:

- ANOVA on CPPs/PPs;
- POUMM/PMM on the whole tree (including tips belonging to CPPs);
- POUMM/PMM on the tree obtained after dropping tips belonging to CPPs;

The results from these analyses are written in table 4. Excluding outlier CPPs, ANOVA-CPP (222 patients) reported lg(spVL)-heritability estimates of 0.31, 95% CI [0.19, 0.43]. POUMM (8,473 patients) reported agreeing estimates of 0.25, 95% CI [0.16, 0.36] and 0.22, CI [0.13, 0.35] upon omitting all 222 patients belonging to CPPs. The slightly lower POUMM estimates could be explained by errors in the transmission tree, which are not present in CPPs. These results show first, that ANOVA-CPP and POUMM agree on disjoint subsets of the UK data and, second, that POUMM provides an alternative to resemblance-based methods in the absence of early-diagnosed cases.

**Table 4.**
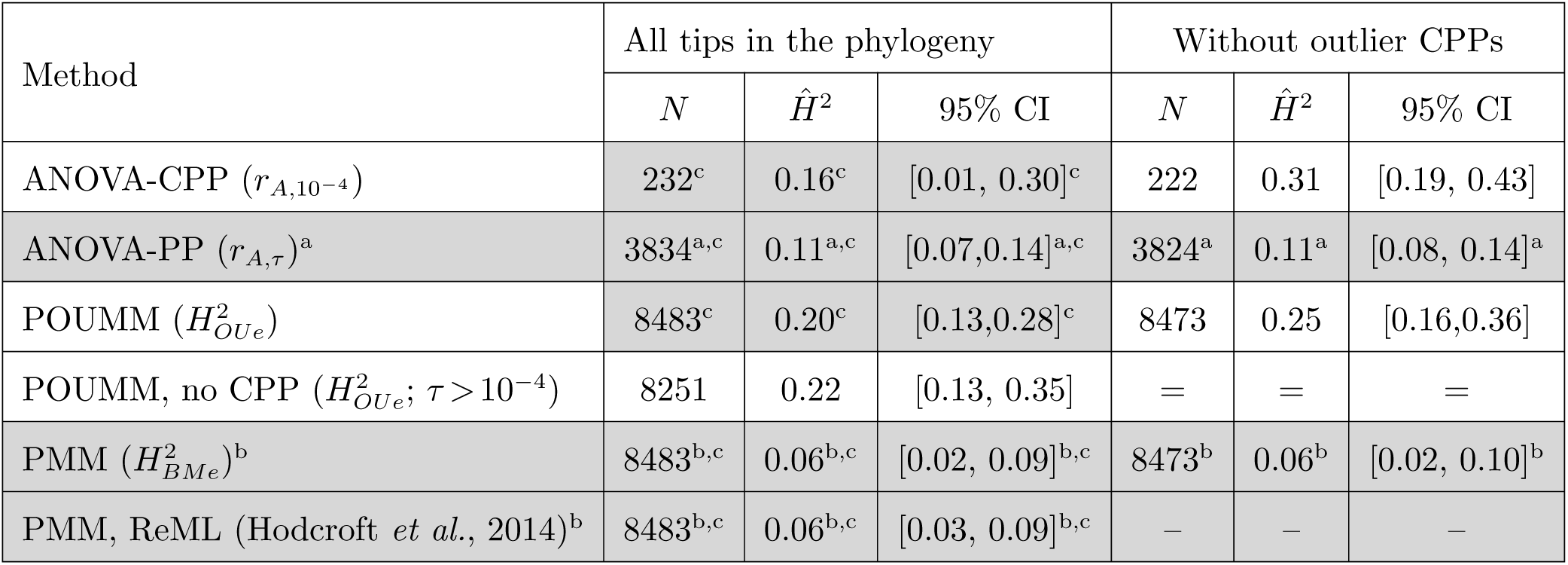
ANOVA-CPP and POUMM estimates of lgspVL-heritability in HIV data from UK. Also written are the results from a previous analysis on the same dataset (Hodcroft et al. 2014). “=”: the input data (and MCMC prior) is not altered by filtering out outlier CPPs; “–”: the analysis was not done in the mentioned study. Grey background: estimates considered unreliable due to: a: negative bias caused measurement delays; b: negative bias caused by BM violation; c: presence of outlier CPPs;

| Method | All tips in the phylogeny |  |  | Without outlier CPPs |  |  |
| --- | --- | --- | --- | --- | --- | --- |
| | $N$ | $\hat{H}^2$ | 95% CI | $N$ | $\hat{H}^2$ | 95% CI |
| ANOVA-CPP ($r_{A,10^{-4}}$) | 232<sup>c</sup> | 0.16<sup>c</sup> | [0.01, 0.30]<sup>c</sup> | 222 | 0.31 | [0.19, 0.43] |
| ANOVA-PP ($r_{A,\tau}$)<sup>a</sup> | 3834<sup>a,c</sup> | 0.11<sup>a,c</sup> | [0.07, 0.14]<sup>a,c</sup> | 3824<sup>a</sup> | 0.11<sup>a</sup> | [0.08, 0.14]<sup>a</sup> |
| POUMM ($H_{OUE}^2$) | 8483<sup>c</sup> | 0.20<sup>c</sup> | [0.13, 0.28]<sup>c</sup> | 8473 | 0.25 | [0.16, 0.36] |
| POUMM, no CPP ($H_{OUE}^2$; $\tau > 10^{-4}$) | 8251 | 0.22 | [0.13, 0.35] | = | = | = |
| PMM ($H_{BMe}^2$)<sup>b</sup> | 8483<sup>b,c</sup> | 0.06<sup>b,c</sup> | [0.02, 0.09]<sup>b,c</sup> | 8473<sup>b</sup> | 0.06<sup>b</sup> | [0.02, 0.10]<sup>b</sup> |
| PMM, ReML (Hodcroft et al., 2014)<sup>b</sup> | 8483<sup>b,c</sup> | 0.06<sup>b,c</sup> | [0.03, 0.09]<sup>b,c</sup> | – | – | – |

Figure 6 compares these estimates to previous lg(spVL) studies using phylogenetic and known transmission-pairs data. In agreement with the toy-model simulations, estimates of *H*^2^ using PMM or other phylogenetic methods (i.e. Blomberg’s *K* and Pagel’s *λ*) are notably lower than all other estimates, suggesting that these phylogenetic comparative methods underestimate *H*^2^; resemblance-based estimates are down-biased by measurement delays (compare early vs late on fig. 6).

**FIG. 6.**
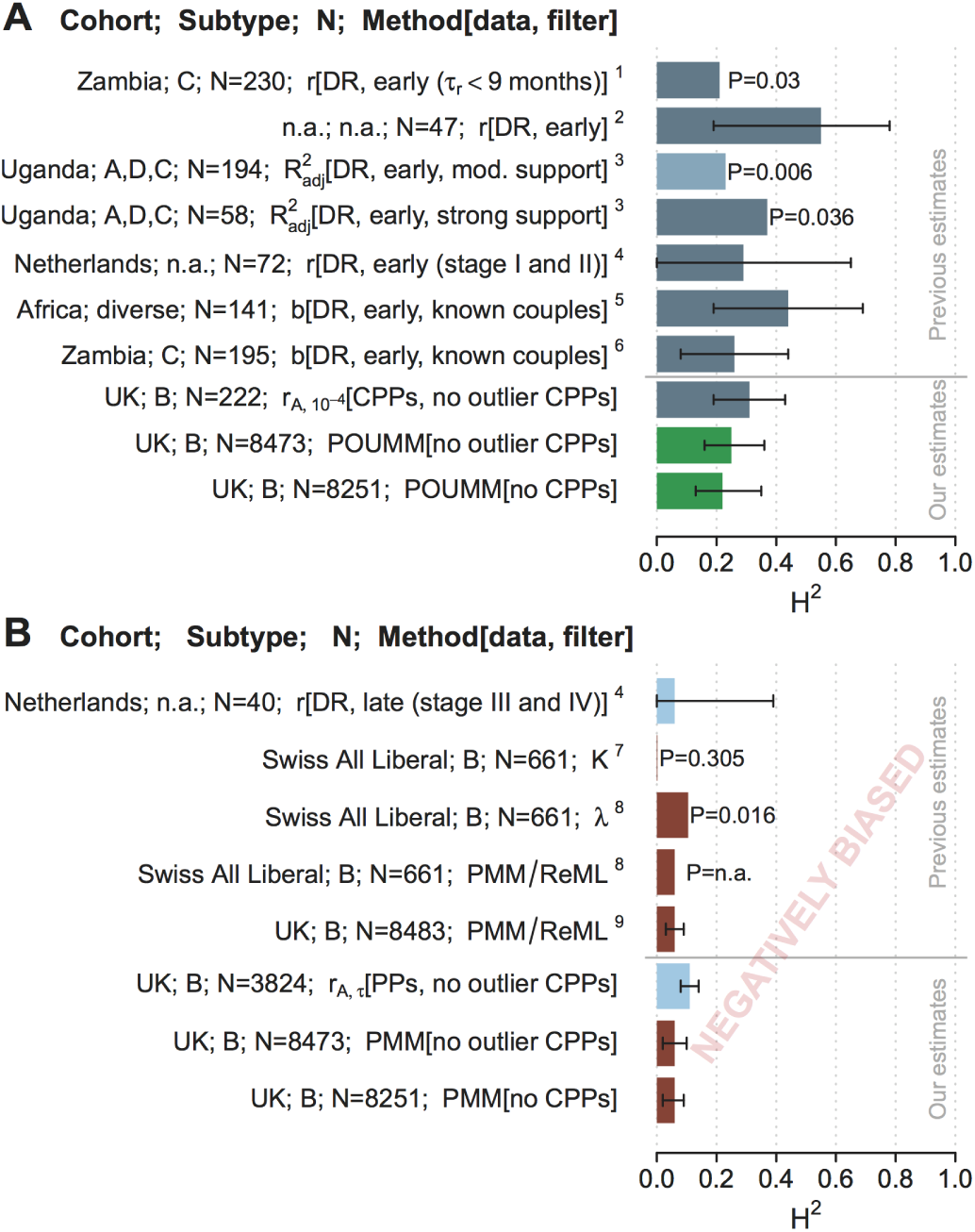
A comparison between *H*^2^-estimates from the UK HIV-cohort and previous estimates on African, Swiss and Dutch data. (A) Estimates with minimized measurement delay (dark cadet-blue) and POUMM estimates (green); (B) Down-biased estimates due to higher measurement delays (light-blue) or violated BM-assumption (brown). Confidence is depicted either as segments indicating estimated 95% CI or P-values in cases of missing 95% CIs. References to the corresponding publications are written as numbers in superscript as follows: 1: Tang et al. 2004; 2: Hecht et al. 2010; 3: Hollingsworth et al. 2010; 4: Van der Kuyl et al. 2010; 5: Lingappa et al. 2013; 6: Yue et al. 2013; 7: Alizon et al. 2010; 8: Shirreff et al. 2013; 9: Hodcroft et al. 2014. For clarity, the figure does not include estimates from the UK data including the five outlier CPPs (table 4) and estimates from previous studies, which are not directly comparable (e.g. previous results from Swiss MSM/strict datasets (Alizon *et al.*, 2010)).

In summary, POUMM and ANOVA-CPP yield agreeing estimates for *H*^2^ in the UK data and these estimates agree with DR-based estimates in datasets with short measurement delay (different African countries and the Netherlands). Similar to the toy-model simulations, we notice a well-pronounced pattern of negative bias for the other estimators, PMM and ANOVA-PP, as well as for the previous DR-studies on data with long measurement delay.

## Discussion

### Clarifying the terminology and notation

The first task of this study was the transfer of quantitative genetics terminology to the domain of pathogen traits. Due to important lifecycle differences between pathogens and mating organisms, it is essential to disentangle the concepts of relative resemblance and genetic determination. In essence, the estimators of trait resemblance between transmission-related patients, such as DR and ICC, and the phylogenetic heritability, must be regarded as lower bounds for the broad-sense heritability, *H*^2^, compromised by partial quasispecies transmission, within-host evolution and various violations of model assumptions (table 3). A few examples from recent studies of HIV demonstrate the need for a careful consideration of these concepts. For example, in (Hodcroft *et al.*, 2014) and (Leventhal and Bonhoeffer, 2016) the authors introduce the PMM/ReML and the DR methods for estimating heritability after a definition of the narrow sense heritability, *h*^2^. This can leave a confusing impression that the reported values are estimates of *h*^2^ rather than *H*^2^, because these methods are popular for estimating narrow-sense heritability for sexual species. As another example, in (Fraser *et al.*, 2014; Shirreff *et al.*, 2013), the authors use the lower-case notation “*h*^2^” to denote estimates of *H*^2^. In fact, there are historical reasons to associate the symbol “*h*^2^” with the regression slope, *b* (Fraser *et al.*, 2014; Wright, 1934). However, “*h*^2^” is the standard symbol for narrow-sense heritability and *b* is, most of all, a measure of phenotypic resemblance. To avoid confusion, we recommend using the standard symbol “*H*^2^” for broad-sense heritability (Hartyl and Clark, 2007; Lynch and Walsh, 1998) and different symbols for its indirect estimators.

### A disagreement between simulation studies

Using simulations of a phenomenological epidemiological model, we have shown that two methods based on phenotypic and sequence data from patients - ANOVA-CPP and POUMM - provide more accurate heritability estimates compared to previous approaches like DR and PMM. However, we should not neglect the arising discrepancy between our and previous simulation reports advocating either PMM (Hodcroft *et al.*, 2014) or DR (Leventhal and Bonhoeffer, 2016) as unbiased heritability estimators. Compared to these simulations, the toy-model presented here has several important advantages: (i) it is biologically motivated by phenomena such as pathogen mutation during infection, transmission of entire pathogens instead of proportions of trait values, and within-/between-host selection; (ii) it is a fair test for all estimators of heritability, because it doesn’t obey any of the estimators’ assumptions, such as linearity of recipient- on donor values, normality of trait values, OU or BM evolution, independence between pathogen and host effects; (iii) it generates transmission trees that reflect the between-host dynamics, e.g. clades with higher trait-values exhibit denser branching in cases of between-host selection. As a criticism, we note that the toy-model does not allow strain coexistence within a host and, thus, is not able to model partial quasispecies transmission and, in particular, transmission bottlenecks (Keele *et al.*, 2008) or preferential transmission of founder strains (Lythgoe and Fraser, 2012). Although it may be exciting from a biological point of view, the inclusion of strain coexistence comes with a series of conceptual challenges, such as the definition of genotype and clonal identity, the formulation of the trait-value as a function of a quasispecies-instead of a single strain genotype, etc. These challenges should be addressed in future studies implementing more advanced models of within-host dynamics and leveraging deep sequencing data. To conclude, the discrepancy between simulation studies teaches that no method suits all simulation setups *ergo* biological contexts. Thus, rather than proving universality of a particular method, simulations should be used primarily to study how particular biologically relevant features affect the methods on table.

### The heritability of HIV set-point viral load is at least 25%

Applied to data from the UK, ANOVA-CPP and POUMM reported four to five times higher point estimates and non-overlapping CIs compared to a previous PMM/ReML-based estimate on the same data (0.06, 95% CI [0.02, 0.09]) (Hodcroft *et al.*, 2014). Our PMM implementation confirmed this estimate. However, based on our simulations (fig. 2 and fig. 4), these estimates are still underestimates of the true heritability. Overall, our analyses yield an unprecedented agreement between estimates of donor-recipient resemblance and phylogenetic heritability in large European datasets and African cohorts, provided that measurements with large delays have been filtered out prior to resemblance evaluation (Hecht *et al.*, 2010; Hollingsworth *et al.*, 2010) (fig. 6A). Also noteworthy are the facts that our estimates for the UK dataset support the results from Fraser *et al.* (2014) who conducted a meta-analysis of three datasets on known transmission partners (Hollingsworth *et al.*, 2010; Lingappa *et al.*, 2013; Yue *et al.*, 2013) (433 pairs in total) reporting heritability values of 0.33, CI [0.20,0.46], as well as the recent results from Blanquart *et al.* (2017) who conducted a POUMM analysis on a whole-genome meta-dataset (1581 sequences from several European countries) reporting spVL heritability of 0.31, CI [0.15, 0.43]. In analogy with our ANOVA approach, Blanquart *et al.* (2017) measured the Pearson correlation in “cherries” partitioned by phylogenetic distance and showing a similar pattern of decreasing correlation (Fig. 2). All datasets support the hypothesis of HIV influencing spVL (*H*^2^*>*0.25). The particular estimates provided in this work should be interpreted as lower bounds for *H*^2^, because the partial quasispecies transmission, the noises in spVL measurements and the noise in transmission trees are included implicitly as environmental (non-transmittable) effects. The non-zero heritability motivates further HIV whole-genome sequencing (Metzner, 2016) and genome-wide studies of the viral genetic association with viral load and virulence.

### A critical view on the POUMM

The OU process has found previous applications as a model for stabilizing selection in macro-evolutionary studies (Felsenstein, 1988; Hansen, 1997; Hansen and Bartoszek, 2012; LANDE, 1976) and references therein. As a contribution of this work, we have shown that the OU process is well adapted for the modeling of pathogen evolution along transmission trees in both, neutral as well as selection scenarios. Unlike BM, OU models the phenotypic resemblance between transmission related patients as a function of their phylogenetic distance, thus, capturing the gradual loss of resemblance caused by within-host evolution (fig. 2). Most of the above-mentioned studies and the accompanying software packages have assumed that the whole trait evolves according to an OU process, usually disregarding the presence of a biologically relevant non-heritable component *e* or treating it as a measurement error whose variance is a priori known (FitzJohn, 2012). Having the OU process act on the genotypic values rather than whole trait-values is a simplifying assumption facilitating mathematical processing (Mitov and Stadler, 2017). However, our toy model simulations have shown robustness and statistical power of the POUMM in complicated scenarios combining trait-based selection at the within- and between-host levels. Another criticism that can be addressed to the POUMM method is that it is unaware of between-host selection and demographic processes, which may result in a correlation between tree structure and trait values (for example higher branching density in clades with higher *z*). As noted by Leventhal and Bonhoeffer (2016), this is a general issue with phylogenetic comparative approaches assuming a global evolutionary process acting on the whole phylogeny. An unexplored alternative would be to associate different instances of POUMM to different clades in the tree based on prior knowledge about heterogeneity between these clades.

### Outlook

ANOVA-CPP and POUMM have great potential to become widely used tools in the study of pathogens. ANOVA-CPP works on pairs of trait values from carriers of nearly identical strains and can be easily extended to groups of variable size (Anderson *et al.*, 2010; Lynch and Walsh, 1998). Thus, ANOVA-CPP is ideal for slowly evolving pathogens such as DNA-viruses, bacteria and protozoa, where clusters of patients carrying identical-by-descent (IBD) strains are frequently found. For example, Anderson et al. 2010 identified 27 clusters of two to eight carriers of IBD strains in a small set of 185 malaria patients, i.e. 41% of the patients participated in clusters (Anderson *et al.*, 2010). On the other hand, IBD-pairs are rare for rapidly evolving RNA-viruses, such as HIV and HCV. For instance, we identified only 116 CPPs in a large dataset of 8483 HIV-sequences, i.e. less than 3% of the patients involved in IBD-pairs. However, the rapidly accumulating sequence diversity of RNA-viruses allows building large-scale phylogenies, which approximate transmission trees between patients. Thus, RNA-viruses should make the ideal scope for the POUMM. We believe that, together, the two methods should enable accurate and robust heritability estimation in a broad range of pathogens.

## Materials and Methods

### Formal definitions of heritability

Here, we briefly review the formal definitions of heritability in sexually reproducing populations based on the general linear model of quantitative traits (Falconer, 1996; Hartyl and Clark, 2007; Lynch and Walsh, 1998) and the three concepts introduced in the main text: the genetic determination of a trait, the resemblance between relatives, and the efficiency of selection.

#### The general linear model of a quantitative trait

A principal goal of quantitative genetics is to partition the observed phenotypic variance in a population into components attributable to genetic and environmental factors. Fundamental for the study of the genetic and environmental sources of variance is the general linear model for the phenotype (see Lynch and Walsh (1998), ch. 6), in which, for a given trait of interest, the observed phenotypic value, *z,* of an organism is represented as a sum of effects of the organism’s genes, *G,* general (macro-) environmental effects, *E,* gene by (macro-) environment interaction, *I,* and special (micro-) environmental effects *e*

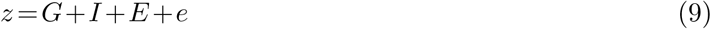

It is assumed that the trait is influenced by a number of genes whose locations in the species’ reference genetic sequence are called quantitative trait loci (QTL). In an individual, the configuration of alleles found at the trait’s QTLs is called genotype and, for a population, the genotypic value, *G***_x_**, of a genotype **x** is defined as the expected trait value of its carriers: *G***_x_** = E(*z|*genotype = **x**). The remaining terms in eq. 9 are “defined in a least-squares sense as deviations from lower order expectations” (Lynch and Walsh, 1998). It is worthy to note that *G***_x_** depends on the distribution of **x** across environments in the population and that, by construction, the residuals *z−G* = *I* +*E* +*e* have zero mean and are uncorrelated with *G* (Lynch and Walsh (1998), ch. 6). Thus, the total phenotypic variance observed in the population can be partitioned into a component that is purely genetic and a component that is attributable to both, non-genetic (purely environmental) factors as well as gene-by-environment interactions: *σ*^2^(*z*) = *σ*^2^(*G*)+*σ*^2^(*z -G*).

#### Measuring the genetic determination of a trait

**Heritability in the broad sense**, a.k.a. degree of genetic determination (Falconer, 1996), is defined as the ratio of the variance of genotypic values to total phenotypic variance in the population:

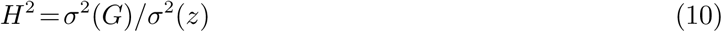

A direct estimation of *H*^2^ would require that all QTLs were known and that for each genotype there was a sample of measurements from individuals who were: (i) genetically identical at the QTLs; (ii) raised in randomly and independently assigned environments; (iii) present in the final dataset according to the population-specific environment-genotype frequencies. Given such a dataset of *N* independent measurements from carriers of all *K* distinct genotypes in the population (*K* ≪ *N*), *H*^2^ can be estimated by the ratio of sample variances 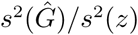, where 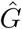 denotes the individuals’ genotypic values estimated by the mean value of their corresponding group and *s*^2^(*·*) denotes sample variance. Though, intuitive, this formula is slightly positively biased in the case of finite sample size. Thus, we prefer its correction for finite degrees of freedom, a.k.a. as adjusted coefficient of determination:

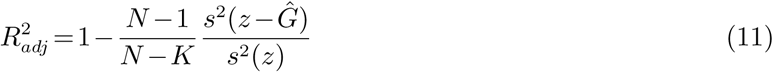

In the absence of full QTL information and data from independently grown clones, direct estimation of *H*^2^ is rarely possible. Instead, quantitative geneticists focus on estimating its lower bound defined below.

**Heritability in the narrow sense** is defined as the ratio of variance of additive genetic values to total phenotypic variance:

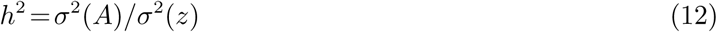

The additive genetic value, *A*, of an organism is defined as the sum of additive effects of its alleles at the trait’s QTLs. We provide the technical definition of additive effect later on and note here that *h*^2^ represents the largest proportion of phenotypic variance that can be explained by linear regression on the allele contents at single QTLs, ignoring epistatic (inter-locus) and dominance interactions (Lynch and Walsh, 1998). As discussed shortly, for sexually reproducing species, *h*^2^ has two main advantages to *H*^2^: (i) it can be estimated from empirical data of genetically related (but not identical) organisms; (ii) it can be used to predict the response to selection for traits associated with reproductive fitness.

#### Measuring the resemblance between relatives

Relatives resemble each other not only for carrying similar sets of alleles but also for living in similar environments. Thus, it is necessary to disentangle the concept of resemblance from that of genetic determination.

Considering an ordered relationship such as parent-offspring, the least squares regression slope of offspring values on mean parental values is defined as

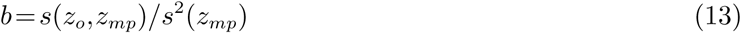

where *z*_*o*_ and *z*_*mp*_ denote observed offspring and mean parent values, and *s*(*·,·*) denotes sample covariance among observed couples of values (Lynch and Walsh, 1998). Assuming no systematic dissimilarity between parents and offspring, *b* is a value between 0 and 1, higher values indicating closer resemblance between the expected phenotype of offspring and the mid-phenotype of their parents.

Considering members of unordered relationships, such as identical twins, sibs and cousins, the resemblance between members within groups is measured by the intraclass correlation (ICC) defined as the ratio of the “between group” variance over the total variance, *r*_*A*_ = *σ*^2^(*c*)/*σ*^2^(*z*), *c* denoting the observed within-group means (Fisher, 1925; Lynch and Walsh, 1998). Given a dataset of measurements grouped by a factor such as twinship, the standard estimation procedure for *r*_*A*_ is the one-way analysis of variance - ANOVA (see, e.g. (Donner, 1986) or ch. 18 in (Lynch and Walsh, 1998)). ANOVA uses mean squares to find estimators for the between- and within-group variances, 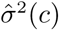 and 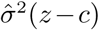 and reports ICC as the ratio:

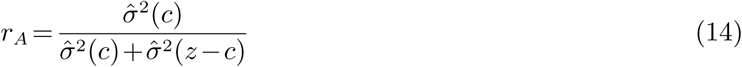

We notice that both, 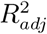 eq. 11) and *r*_*A*_ (eq. 14), are estimators of ICC, but there is a key difference in their assumptions: 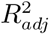assumes that all possible groups, i.e. genotypes, are present in the data but makes no explicit assumption about the distribution of group means (i.e. genotypic values); *r*_*A*_ is aware that only a subset of all possible groups is present in the data but assumes that the observed group means, are an i.i.d. sample from a normal distribution.

#### Measuring the efficiency of selection

In breeding experiments the goal is to optimize a trait by repetitive artificial selection for reproduction of the “best” individuals in a generation. A textbook example is truncation selection in which only individuals with measurements above a given threshold are allowed to reproduce. For a generation, the difference Δ_*s*_ = *μ*_*s*_ *−μ* between the mean value of individuals selected for reproduction, *μ*_*s*_, and the mean of the generation, *μ*, is called the selection differential. Denoting by the mean of the offspring generation, the difference *R* = *μ*_*o*_ *−μ*, is called the response to selection. Then, the efficiency of the truncation selection is measured by the **realized heritability** (Hartyl and Clark, 2007), defined as the ratio:

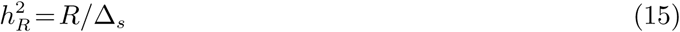

#### Definition of additive genetic effect and additive genetic value

So far, we have skipped the more technical definition of additive genetic effect, which is the basis of the definitions of additive genetic value and narrow-sense heritability. Here we provide these definitions in the context of haploid organisms, noting that the definitions for diploid organisms found in textbooks (Falconer, 1996; Lynch and Walsh, 1998) are conceptually the same but somewhat more complicated for they treat dominance interactions separately from epistatic interactions.

We assume that a trait has a finite number of QTLs, *L*, with a finite number of alleles *M*_*l*_ ⩾ 2 for each locus *l* = 1*,…,L*. Denoting by *x*_*lm*_ the content (0 or 1) of allele *m* at locus *l*, *l* = 1*,…,L*, *m* = 1*,…,M*_*l*_, we can describe an individual’s genotype by a binary vector **x** of length 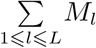. The products of allele contents for different loci signify the presence or absence of allele combinations in a genotype. This representation results in the system of equations 16, in which the genotypic value of each genotype **x** is written as a sum of the population mean, *μ*, and the effects *η*_*lm*_, (*ηη*)_*l*_1_ *m*_1_ *l*_2_ *m*_2__ and so on, associated with each allele, couple of alleles at two loci and higher order- (up to order *L*) multi-locus configurations of alleles, present in the genotype.

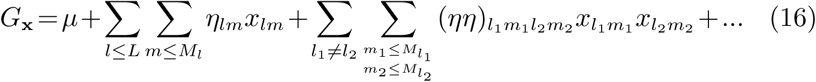

If for a moment we imagine that in system of equations *G***_x_**, *μ*, and **x** are known while the (*η…*)′**s** are unknown, from an algebraic point of view, there exist infinitely many combinations of (*η…*)^′^**s** solving the system, because there are more unknowns than equations. From the point of view of genetics, however, useful solutions are only those that maximize the proportion of variance in the genotypic values explained by the effects of single alleles or low-order allele combinations. This reasoning finds a mathematical reflection in the ordinary least squares (OLS) solution for the linear regression of *G***_x_** on single-locus allele contents **x** (system 16 taken without the grey-shaded higher order terms on the right). Denoting by *f***_x_** the frequency of genotype **x** among individuals in the population, the vector of OLS coefficients, *η\**, is found as a solution to the optimization task 17:

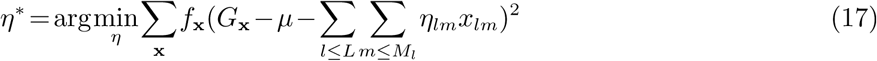

The elements 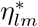 of any vector *η\** solving this optimization task are called **additive allele effects** and the sum 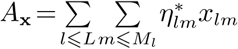 is called **additive genetic value** of the genotype **x**. As a detail, we clarify that for multiple QTLs (*L>*1) the vector *η\** solving 17 is not uniquely defined because for each locus one of the allele contents can be expressed as a function of the others, i.e. the design matrix of the linear model is not of full rank. However the additive genetic values are invariant to the exact choice of *η\**.

### Software

This study relies on the accompanying R-package “patherit”. The used version of this package, together with all program-code used for the toy-model simulations and the analysis of HIV-data, are provided in the attached file SP.zip. Inside it, a file named ReadMe.txt contains further instructions on how to run the code. The sub-sections below provide details on the implementation of the different heritability estimators and the toy-model simulations.

#### Direct measurement of H^2^ in simulated data

To measure *H*^2^, we used the direct estimate 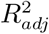 (Eq. 11) after grouping the patients in the data by their (currently carried) pathogen genotype and estimating the genotypic values as the group means (implemented as function R2adj in the patherit package).

#### Calculating donor-recipient regression slope

The value of the donor-recipient regression slope (*b*_0_, *b*_*D*_1__, *b*_*τ*_) was calculated using eq. 13, implemented as a function called “b” in the patherit package.

#### Calculating r_A_

To estimate *r*_*A*_ we implemented one-way ANOVA as a function “rA” in the package patherit. As a reference we used the description in chapter 18 of (Lynch and Walsh, 1998). To calculate confidence intervals, we used the R-package “boot” to perform 1,000-replicate bootstraps, upon which we called the package function boot.ci() with type=”basic”. These confidence intervals were fully contained in the standard ANOVA confidence intervals based on the F-distribution (see (Lynch and Walsh, 1998)), which were slightly wider (not reported).

#### POUMM and PMM inference

The POUMM and PMM inference was based on an early version of the POUMM R-package (Mitov and Stadler, 2017). Since the interface of the POUMM package has evolved considerably between the version used in this analysis and the version 1.2.1 released on the Comprehensive R Archive Network (CRAN, https://cran.r-project.org) at the time of writing this article. To facilitate reproducibility, the source-code of the early version used in this analysis has been included in the accompanying package ’patherit’.

We performed maximum likelihood (ML) fits of the POUMM method in all toy model simulations. For each simulated transmission tree, the conditional likelihood of the trait-values at the tips was maximized over the parameters *α*, *θ*, *σ*, *σ*_*e*_ and *g*_0_ (function ml.poumm of the patherit package). In the PMM ML fits the conditional likelihood of the data was redefined as its corresponding limit for *α →* 0 and was maximized over the parameters *σ*, *σ*_*e*_ and *g*_0_ (ignoring *θ*, which cancels out in the case *α →* 0). To avoid potential issues with floating point arithmetic all branch lengths were scaled-down 100 times before ML fit. This preprocessing step is invariant with respect to the estimated heritability, since it only causes rescaling of the OU parameters: *σ*^2^ *→ σ*^2^ *×*100; *α → α ×*100 (see eq. 5).

For HIV data, in addition to an ML-fit, we performed a Markov Chain Monte Carlo (MCMC) fit (function mcmc.poumm of the patherit package) using an adaptive Metropolis algorithm with coerced acceptance rate (Vihola, 2012) written in R (Scheidegger, 2012). The MCMC sampling was performed on the POUMM parameters *α*, *θ*, *σ*^2^ and 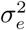. The prior was specified as a joint distribution of four independent variables: 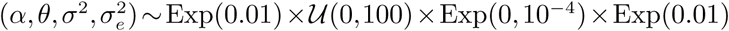. These low exponential rates and the large interval of the uniform distribution were chosen such in order to ensure that the prior is weakly informed, both, for the sampled parameters *α*, *θ*, *σ*^2^, 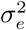 and for the inferred heritability estimates 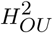,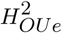. This is verified by the nearly flat prior densities contrasting with sharply peaked posterior densities (compare blue versus black curves on supplementary fig. S4 B). The initial values for the parameters were set to 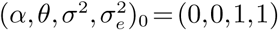. The adaptive Metropolis MCMC was run for 4.2E+06 iterations, of which the first 2E+05 were used for warm-up and adaptation of the jump distribution variance-covariance matrix. The target acceptance rate was set to 0.01 and the thinning interval was set to 1,000. The convergence and mixing of the MCMC was validated by visual analysis (supplementary fig. S4 A) as well as by comparison to a parallel MCMC-chain started from a different initial state. The presence of signal in the data was confirmed by the observed difference between prior (blue) and posterior (black) densities (see supplementary fig. S4 B). Calculation of 95% CI was done using the function “HPDinterval” from the coda package (Plummer *et al.*, 2006).

#### Computer simulations of the toy epidemiological model

The toy-model SIR simulation is implemented in the function “simulateEpidemic” of the patherit package; the extraction of diagnosed donor-recipient couples – in the function “extractDRCouples”; the extraction of a transmission tree from diagnosed individuals – in the function “extractTree”.

At the between-host level, the phenomena of birth, contact, transmission, recovery and death define the dynamics between the compartments of susceptibles, infected and recovered individuals - *X*, *Y* and *Z*. The natural birth rate, *λ*_nat_, and the natural per capita death rate, *δ*_nat_, are defined as constants satisfying *λ*_nat_ = *δ*_nat_*N*_0_, so that the average lifespan of an uninfected individual equals 1*/δ*_nat_ = 850 (arbitrary) time units and in a disease-free population the total number of alive individuals equilibrates at *N*_0_ = 10^5^. An epidemic starts with the migration of an individual with random immune system type carrying pathogen strain 1:11 to a fully susceptible population of *N*_0_ individuals. Each individual has contacts with other individuals occurring randomly at a constant rate, *κ*. A transmission can occur upon a contact involving an infected and a susceptible individual, here, called a “risky” contact. It is assumed that the probability of transmission per risky contact, *γ*, is either a constant (black on fig. 3B) or a function of the value *z* (magenta on Fig. 3B) of the infected host and does not depend on the uninfected individual. Once infected, a host starts transmitting its currently dominant pathogen strain at a rate defined as the product of *γ*, *κ*, and the current proportion of susceptible individuals in the population, *S* = *X/N*. Thus, for fixed *κ*, the transmission rate of an infected host is a function of the global variable *S* and the constant or variable *γ*. This transmission process continues until recovery or death of the host. Recovery has the meaning of a medical check occurring at a constant per capita rate, *ρ*, followed by immediate therapy and immunity. Due to the virulence of the pathogen, an infected host has an increased (per capita) death rate, *δ*, which is defined either as a constant or as a function of *z*. Based on their scope of action, we call “between-host” the parameters *λ*_nat_, *δ*_nat_, *κ*, *γ*, *ρ*and *δ*.

Within a host, mutants of the dominant strain can appear at any time as a result of random single-locus mutations, which occur at a constant or *z*-dependent rate, *ν*. It is important to make a distinction between a mutation and a substitution of a mutant strain for a dominant strain within a host, because a mutation doesn’t necessarily lead to a substitution. For example, when *z* is (or correlates with) the within-host reproductive fitness of the pathogen, substitutions would result only from mutations causing an increase in *z*. The rate of substitution of a mutant strain **x**_*j*_ for a dominant strain **x**_*i*_, differing by a single nucleotide at a locus *l*, is denoted *ξ*_*l,i←j*_ and defined as a function of *ν*, the number of alleles at the locus, *M*_*l*_, and the presence or absence of within-host selection with respect to *z*. No substitution can occur between strains differing at more than one locus, although, the same effect can result from two or more consecutive substitutions. Based on their scope of action, we call “within-host” the parameters *v* and *ξ*.

The parameters *λ*_nat_, *δ*_nat_, *κ* and *ρ* were kept as global constants as written in table 1.

The simulations were implemented as stochastic random sampling of within- and between-host events (i.e. risky contact, transmission, mutation, diagnosis, death) in discrete time-steps of length 0.05 (arbitrary time-units). Each simulation was run for min(4*t*_10*k*_,2400) time-units, where *t*_10*k*_ denotes the time for the simulation until reaching 10,000 diagnosed individuals. The data generated after reaching 10,000 diagnoses has not been used in this study but it is intended for future analysis of post-outbreak dynamics, i.e. epidemic waves occurring after exhaustion of the susceptible pool. The transmission history as well as the history of within-host strain substitutions was preserved during the simulations in order to reproduce exact transmission trees and to extract donor and recipient values at moments of transmission for the calculation of *b*_0_.

#### External dependencies

The following third-party R-packages were used: ape v3.4 (Paradis *et al.*, 2004), data.table v1.9.6 (Dowle *et al.*, 2014), adaptMCMC v1.1 (Scheidegger, 2012), Rmpfr v0.6-0 (Maechler, 2014), and coda v0.18-1 (Plummer *et al.*, 2006). All programs have been run on R v3.2.4 (R Core Team, 2013).

## Supplementary Material

Supplementary notes, figures S1-S4 and supplementary programs are available online.

## Acknowledgments

This work was supported by the Eidgenssische Technische Hochschule Zrich and in part by the European Research Council under the 7th Framework Programme of the European Commission (PhyPD: Grant Agreement Number 335529).

The authors thank Dr. Emma Hodcroft for sending the UK phylogeny in Newick format together with the associated spVL values, Dr. Gabriel Leventhal and prof. Sebastian Bonhoeffer for valuable insights on donor-recipient regression, Dr. Francois Blanquart and prof. Christoph Fraser for sharing with us their early results on the Beehive dataset and for valuable discussions, and Dr. David Rasmussen for a careful review of the manuscript.

## References

Alizon, S., von Wyl, V., Stadler, T., Kouyos, R. D., Yerly, S., Hirschel, B., Böni, J., Shah, C., Klimkait, T., Furrer, H., Rauch, A., Vernazza, P. L., Bernasconi, E., Battegay, M., Bürgisser, P., Telenti, A., Günthard, H. F., Bonhoeffer, S., and Swiss HIV Cohort Study 2010. Phylogenetic approach reveals that virus genotype largely determines HIV set-point viral load. PLoS pathogens, 6(9): e1001123.

Anderson, T. J. C., Williams, J. T., Nair, S., Sudimack, D., Barends, M., Jaidee, A., Price, R. N., and Nosten, F. 2010. Inferred relatedness and heritability in malaria parasites. Proceedings of the Royal Society B-Biological Sciences, 277(1693): 2531–2540.

Bjorn-Mortensen, K., Soborg, B., Koch, A., Ladefoged, K., Merker, M., Lillebaek, T., Andersen, A. B., Niemann, S., and Kohl, T. A. 2016. Tracing Mycobacterium tuberculosis transmission by whole genome sequencing in a high incidence setting: a retrospective population-based study in East Greenland. Scientific reports, 6(1): 33180.

Blanquart, F., Wymant, C., Cornelissen, M., Gall, A., Bakker, M., Bezemer, D., Hall, M., Hillebregt, M., Ong, S. H., Albert, J., Bannert, N., Fellay, J., Fransen, K., Gourlay, A., Grabowski, M. K., Gunsenheimer-Bartmeyer, B., Günthard, H. F., Kivelä, P., Kouyos, R., Laeyendecker, O., Liitsola, K., Meyer, L., Porter, K., Ristola, M., van Sighem, A., Vanham, G., Berkhout, B., Kellam, P., Reiss, P., and Fraser, C. 2017. Viral genetic variation accounts for a third of variability in HIV-1 set-point viral load in Europe. Plos Biology, In press (personal communication).

Bonhoeffer, S., Fraser, C., and Leventhal, G. E. 2015. High Heritability Is Compatible with the Broad Distribution of Set Point Viral Load in HIV Carriers. PLoS pathogens, 11(2): e1004634–e1004634.

Dessau, M., Goldhill, D., McBride, R. L., Turner, P. E., and Modis, Y. 2012. Selective Pressure Causes an RNA Virus to Trade Reproductive Fitness for Increased Structural and Thermal Stability of a Viral Enzyme. PLOS Genetics, 8(11).

Donner, A. 1986. A Review of Inference Procedures for the Intraclass Correlation Coefficient in the One-Way Random Effects Model. International Statistical Review, 54(1): 67–82.

Dowle, M., Short, T., Liangolou, S., and Srinivasan, A. 2014. data.table: Extension of data.frame. page 9.

Falconer, D. S. 1996. Introduction to Quantitive Genetics. San Val, Incorporated.

Felsenstein, J. 1988. Phylogenies And Quantitative Characters. Annual Review of Ecology and Systematics, 19(1): 445–471.

Fisher, R. A. 1925. Statistical Methods For Research Workers. Genesis Publishing Pvt Ltd.

FitzJohn, R. G. 2012. Diversitree: comparative phylogenetic analyses of diversification in R. Methods in Ecology and Evolution, 3(6): 1084–1092.

Fraser, C., Hollingsworth, T. D., Chapman, R., de Wolf, F., and Hanage, W. P. 2007. Variation in HIV-1 set-point viral load: epidemiological analysis and an evolutionary hypothesis. Proceedings of the National Academy of Sciences of the United States of America, 104(44): 17441–17446.

Fraser, C., Lythgoe, K., Leventhal, G. E., Shirreff, G., Hollingsworth, T. D., Alizon, S., and Bonhoeffer, S. 2014. Virulence and Pathogenesis of HIV-1 Infection: An Evolutionary Perspective. Science (New York, N.Y.), 343(6177): 1243727–1243727.

Freckleton, R. P., Harvey, P. H., and Pagel, M. 2002. Phylogenetic analysis and comparative data: a test and review of evidence. The American Naturalist, 160(6): 712–726.

Geskus, R. B., Prins, M., Hubert, J.-B., Miedema, F., Berkhout, B., Rouzioux, C., Delfraissy, J.-F., and Meyer, L. 2007. The HIV RNA setpoint theory revisited. Retrovirology, 4(1): 65.

Grimmett, G. and Stirzaker, D. 2001. Probability and Random Processes. Oxford University Press.

Hansen, T. F. 1997. Stabilizing Selection and the Comparative Analysis of Adaptation. Evolution; international journal of organic evolution, 51(5): 1341–1351.

Hansen, T. F. and Bartoszek, K. 2012. Interpreting the evolutionary regression: the interplay between observational and biological errors in phylogenetic comparative studies. bioRxiv, 61(3): 413–425.

Hartyl, D. L. and Clark, A. G. 2007. Principles of population genetics. Sinauer Associates.

Hecht, F. M., Hartogensis, W., Bragg, L., Bacchetti, P., Atchison, R., Grant, R., Barbour, J., and Deeks, S. G. 2010. HIV RNA level in early infection is predicted by viral load in the transmission source. AIDS (London, England), 24(7): 941–945.

Hodcroft, E., Hadfield, J. D., Fearnhill, E., Phillips, A., Dunn, D., O'Shea, S., Pillay, D., Brown, A. J. L., Database, o. b. o. t. U. H. D. R., and Study, t. U. C. 2014. The Contribution of Viral Genotype to Plasma Viral Set-Point in HIV Infection. PLoS pathogens, 10(5): e1004112.

Hollingsworth, T. D., Laeyendecker, O., Shirreff, G., Donnelly, C. A., Serwadda, D., Wawer, M. J., Kiwanuka, N., Nalugoda, F., Collinson-Streng, A., Ssempijja, V., Hanage, W. P., Quinn, T. C., Gray, R. H., and Fraser, C. 2010. HIV-1 transmitting couples have similar viral load set-points in Rakai, Uganda. PLoS pathogens, 6(5): e1000876.

Housworth, E. A., Martins, E. P., and Lynch, M. 2004. The phylogenetic mixed model. The American Naturalist, 163(1): 84–96.

Hu, S., Clewley, J. P., Cane, P. A., and Pillay, D. 2004. HIV-1 pol gene variation is sufficient for reconstruction of transmissions in the era of antiretroviral therapy. AIDS (London, England), 18(5): 719–728.

Jacquard, A. 1983. Heritability: One Word, Three Concepts. Biometrics, 39(2): 465.

Keele, B. F., Giorgi, E. E., Salazar-Gonzalez, J. F., Decker, J. M., Pham, K. T., Salazar, M. G., Sun, C., Grayson, T., Wang, S., Li, H., Wei, X., Jiang, C., Kirchherr, J. L., Gao, F., Anderson, J. A., Ping, L.-H., Swanstrom, R., Tomaras, G. D., Blattner, W. A., Goepfert, P. A., Kilby, J. M., Saag, M. S., Delwart, E. L., Busch, M. P., Cohen, M. S., Montefiori, D. C., Haynes, B. F., Gaschen, B., Athreya, G. S., Lee, H. Y., Wood, N., Seoighe, C., Perelson, A. S., Bhattacharya, T., Korber, B. T., Hahn, B. H., and Shaw, G. M. 2008. Identification and Characterization of Transmitted and Early Founder Virus Envelopes in Primary HIV-1 Infection. Proceedings of the National Academy of Sciences of the United States of America, 105(21): 7552–7557.

Keeling, M. J. and Rohani, P. 2007. Modeling Infectious Diseases in Humans and Animals. Princeton University Press.

Lande, R. 1976. Natural-Selection and Random Genetic Drift in Phenotypic Evolution. Evolution; international journal of organic evolution, 30(2): 314–334.

Leventhal, G. E. and Bonhoeffer, S. 2016. Potential Pitfalls in Estimating Viral Load Heritability. Trends in microbiology.

Lingappa, J. R., Thomas, K. K., Hughes, J. P., Baeten, J. M., Wald, A., Farquhar, C., de Bruyn, G., Fife, K. H., Campbell, M. S., Kapiga, S., Mullins, J. I., and Connie Celum, f. t. P. i. P. H. H. T. S. T. 2013. Partner Characteristics Predicting HIV-1 Set Point in Sexually Acquired HIV-1 Among African Seroconverters. http://www.liebertpub.com/, 29(1): 164–171.

Lynch, M. 1991. Methods for the Analysis of Comparative Data in Evolutionary Biology. Evolution; international journal of organic evolution, 45(5): 1065–1080.

Lynch, M. and Walsh, B. 1998. Genetics and Analysis of Quantitative Traits. Sinauer Associates Incorporated.

Lythgoe, K. A. and Fraser, C. 2012. New insights into the evolutionary rate of HIV-1 at the within-host and epidemiological levels. Proceedings: Biological Sciences, 279(1741): 3367–3375.

Maechler, M. 2014. Rmpfr: R MPFR - Multiple Precision Floating-Point Reliable.

Mellors, J. W., Rinaldo, C. R., Gupta, P., White, R. M., Todd, J. A., and Kingsley, L. A. 1996. Prognosis in HIV-1 infection predicted by the quantity of virus in plasma. Science (New York, N.Y.), 272(5265): 1167–1170.

Metzner, K. J. 2016. HIV Whole-Genome Sequencing Now: Answering Still-Open Questions. Journal of clinical microbiology, 54(4): 834–835.

Mitov, V. and Stadler, T. 2017. POUMM: An R-package for Bayesian Inference of Phylogenetic Heritability.

Paradis, E., Claude, J., and Strimmer, K. 2004. APE: Analyses of Phylogenetics and Evolution in R language. Bioinformatics, 20(2): 289–290.

Plummer, M., Best, N., Cowles, K., and Vines, K. 2006. CODA: Convergence Diagnosis and Output Analysis for MCMC. R News, 6: 7–11.

Presloid, J. B., Mohammad, T. F., Lauring, A. S., and Novella, I. S. 2016. Antigenic diversification is correlated with increased thermostability in a mammalian virus. Virology, 496: 203–214.

R Core Team 2013. R: A Language and Environment for Statistical Computing.

Scheidegger, A. 2012. adaptMCMC: Implementation of a generic adaptive Monte Carlo Markov Chain sampler.

Shirreff, G., Alizon, S., Cori, A., Günthard, H. F., Laeyendecker, O., van Sighem, A., Bezemer, D., and Fraser, C. 2013. How effectively can HIV phylogenies be used to measure heritability? Evolution, Medicine, and Public Health, 2013(1): 209–224.

Tang, J., Tang, S., Lobashevsky, E., Zulu, I., Aldrovandi, G., Allen, S., Kaslow, R. A., and Zambia-UAB HIV Research Project 2004. HLA allele sharing and HIV type 1 viremia in seroconverting Zambians with known transmitting partners. AIDS research and human retroviruses, 20(1): 19–25.

Uhlenbeck, G. E. and Ornstein, L. S. 1930. On the Theory of the Brownian Motion. Physical Review, 36(5): 823–841.

van der Kuyl, A. C., Jurriaans, S., Pollakis, G., Bakker, M., and Cornelissen, M. 2010. HIV RNA levels in transmission sources only weakly predict plasma viral load in recipients. AIDS (London, England), 24(10): 1607–1608.

Vihola, M. 2012. Robust adaptive Metropolis algorithm with coerced acceptance rate. Statistics and Computing, 22(5): 997–1008.

Virgin, H. W., Wherry, E. J., and Ahmed, R. 2009. Redefining Chronic Viral Infection. Cell, 138(1): 30–50.

Wright, S. 1934. The Method of Path Coefficients. The Annals of Mathematical Statistics, 5(3): 161–215.

Yue, L., Prentice, H. A., Farmer, P., Song, W., He, D., Lakhi, S., Goepfert, P., Gilmour, J., Allen, S., Tang, J., Kaslow, R. A., and Hunter, E. 2013. Cumulative impact of host and viral factors on HIV-1 viral-load control during early infection. Journal of Virology, 87(2): 708–715.

